# SMN-primed ribosomes modulate the translation of transcripts related to Spinal Muscular Atrophy

**DOI:** 10.1101/751701

**Authors:** F Lauria, P Bernabò, T Tebaldi, EJN Groen, E Perenthaler, M Clamer, F Maniscalco, M Marchioretto, Julia Orri, M Dalla Serra, A Inga, A Quattrone, TH Gillingwater, G Viero

**Affiliations:** Institute of Biophysics, CNR Unit at Trento, Italy; Department CIBIO, University of Trento, Trento, Italy; Edinburgh Medical School: Biomedical Sciences & Euan MacDonald Centre for Motor Neurone Disease Research, University of Edinburgh, Edinburgh, UK; Immagina Biotechnology s.r.p., Trento (Italy); Yale Comprehensive Cancer Center, Yale University School of Medicine, New Haven, CT, USA; Department of Neurology and Neurosurgery, UMC Utrecht Brain Center, Utrecht, the Netherlands; Department of Clinical Genetics, Erasmus University Medical Center, Rotterdam, The Netherlands

## Abstract

The contribution of ribosome heterogeneity and ribosome-associated factors to the molecular control of proteomes in health and disease remains enigmatic. We demonstrate that Survival Motor Neuron (SMN) protein, loss of which causes the neuromuscular disease spinal muscular atrophy (SMA), binds to ribosomes and that this interaction is tissue-dependent. SMN-primed ribosomes are positioned within the first five codons of a set of mRNAs which are enriched in IRES-like sequences in the 5’UTR and rare codons at the beginning of their coding sequence. Loss of SMN at early-stages of SMA induces translational defects *in vivo,* characterized by ribosome depletion in rare codons at the third and fifth position of the coding sequence. These positional defects cause ribosome depletion from mRNAs bound by SMN-primed ribosomes and translational impairment of proteins involved in motor neuron function and stability, including acetylcholinesterase. Thus, SMN plays a crucial role in the regulation of ribosome fluxes along mRNAs which encode proteins relevant to SMA pathogenesis.

## Introduction

As soon as eukaryotic mRNA is transcribed and spliced it is exported to the cytoplasm. Here, it is examined, modified and transported by, or stored in, RNA granules, before eventually being translated into proteins by ribosomes in polysomes. Translation is the most energy consuming process in cells (*1*, *2*) and represents a core mechanism coordinating multiple post-transcriptional processes. Hence, it is not surprising that several mRNAs are largely controlled at the translational rather than transcriptional level (*3–5*). Indeed, loss of post-transcriptional and translational control has been linked to cancer (*6*, *7*), autism (*8*) and neurodegenerative disease (*9–11*), highlighting the critical contribution of translation to disease pathogenesis.

Ribosomes have been placed in the spotlight as putative direct players in tuning translation by acting as mRNA regulatory elements, or “filters” (*12*, *13*). Recent findings also suggest that ribosome composition is not fixed and uniform, but rather is heterogeneous and can be modulated at both the level of ribosomal protein composition (*14–16*), rRNA variants (*17*, *18*) or by ribosome-associated factors (RAFs) (*12*, *19*). Indeed, ribosome heterogeneity is considered to exert a direct role on mRNA selection (*15*) and function (*19*, *20*). Although this represents an exciting potential mechanism for ribosome-based control of gene expression, at present it remains unclear whether direct or indirect defects in ribosome heterogeneity can contribute to disease pathogenesis.

Loss of the Survival Motor Neuron protein (SMN), by homozygous deletion or genetic mutations in *SMN1*, causes spinal muscular atrophy (SMA) (*21*, *22*). SMA is characterized by loss of lower motor neurons, leading to muscle atrophy and wasting. However, the molecular mechanisms leading to motor neuron death in SMA remain complex and unresolved (*22–26*). Although classically known to play a role in the biogenesis of ribonucleoparticles (RNPs) (*27*), SMN protein is a strong candidate to be directly implicated in the control of translation. In fact, SMN is thought to associate with polysomes in cell cultures (*11*, *28*), as well as rat and mouse spinal cords (*11*, *29*) and mouse brain (*11*). Moreover, SMN influences translation both *in vitro* (*11*, *28*, *30*) and *in vivo* (*11*). Hence, it is possible that, in addition to its known roles (*27*), SMN protein functions as a ribosome modulator leading to early and local dysfunction of translation when levels of SMN are decreased. In line with this hypothesis, genome-wide defects occurring in mRNA recruitment onto polysomes have previously been observed in SMA (*11*), but the mechanism(s) linking SMN to these defects in SMA have yet to be elucidated.

Here, we present evidence suggesting that SMN is a ribosome-associated factor acting as a master regulator of translation on a specific subset of mRNAs relevant to SMA pathogenesis.

## Results

### SMN binds ribosomes in vitro and in vivo

Guided by previous evidence suggesting an association of SMN with polysomes (*11*, *28–30*), we hypothesised that SMN may play an as yet uncharacterized role in regulating translation by acting as a ribosome-associated factor. To detail the interaction between SMN and ribosomes, we first characterized the binding of recombinant SMN to purified SMN-free ribosomes obtained from cells which do not express SMN (*11*) (**Figure 1A**). We found that recombinant SMN binds isolated ribosomes *in vitro* with Kd = 180 nM (**Figure 1B**, **Supplementary Figure 1A**). Next, we performed a sub-cellular fractionation coupled to high salt wash (*31*) (**Figure 1C**) in different mouse tissues. Before and after salt washes, SMN remained tightly associated with ribosomes in both brain (**Figure 1D**) and spinal cord (**Supplementary Figure 1B**). To monitor if the interaction of SMN with ribosomes/polysomes is mRNA dependent, we treated the ribosome/polysome pellet with RNase I and observed that SMN still sediments with ribosomes/polysomes, suggesting that this association is mRNA-independent (**Figure 1E**, **Supplementary Figure 1C**).

**Figure 1.**
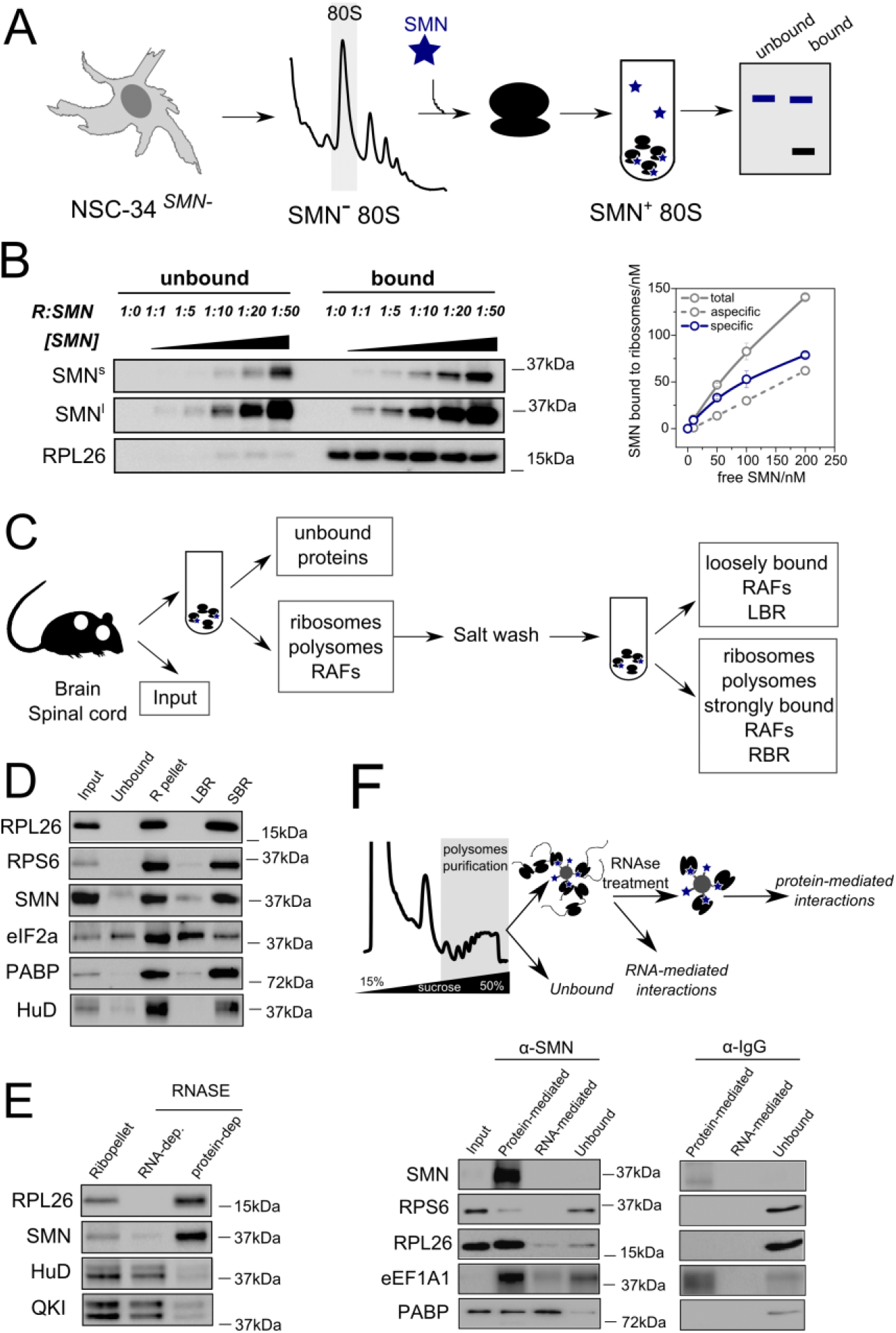
SMN interacts with the translation machinery *in vitro* and *in vivo* in an RNA-independent manner. (**A**) Schematic overview of experimental design for studying the binding of recombinant SMN to purified SMN-free ribosomes. Ribosomes were obtained from SMN-free cell lysates using NSC-34 cells depleted of SMN using CRISPR-Cas9 (*11*). After RNAse digestion, 80S ribosomes were isolated using sucrose gradient fractionation and subsequent ultracentrifugation. Resuspended ribosomes were incubated with recombinant SMN. Bound and unbound SMN were separated by ultracentrifugation and analysed by western blotting. (**B**) Western blotting of unbound and ribosome-associated SMN at different SMN concentrations (10 nM to 500 nM). The concentration of ribosomes was constant (10 nM) in all experiments. Samples were ultracentrifuged to isolate the ribosome pellet and the presence of SMN was determined by western blot. The saturation curve was obtained and the Kd calculated. Experiments were performed using three independent replicates. (**C**) Schematic overview of the subcellular fractionation protocol used to study the association of SMN to ribosomes in brain and spinal cord. (**D**) Western blot analysis on P7 cytoplasmic lysates from brain; input; ribosome-free cytoplasmic components (unbound); ribosomal subunits, ribosomes and polysomes (R-pellet); loosely ribosome-bound proteins (LBR); and strongly ribosome-bound proteins (SBR) from P7 brain. PABP and eIF2a are proteins associated to polysomes and HuD is an RNA binding protein known to be associated to polysomes through RNA interactions. The ribosomal proteins L26 and S6 were used as control of ribosome sedimentation. (**E**) The ribo-pellet was treated with RNase I and ultracentrifuged to separate proteins interacting with ribosomes through RNA-dependent or independent interactions (C). The RNA binding proteins HuD and QKI were used as controls for RNA-dependent interactions with polysomes. (**F**) Scheme of immunoprecipitation of SMN from purified polysomal fractions (upper panel). The fist wash corresponds to the “unbound” lane. After on beads RNase treatment proteins were extracted from beads (Protein-mediated) or from washes (RNA-mediated). The IP was performed on sucrose fractions corresponding to polysomes in brain P5 with anti-SMN or mouse IgG as control (lower panel). Ribosomal proteins L26 and S6 and the elongation factor 1A1 interact via protein-dependent interactions and Poly-A Binding Protein via a protein-dependent or RNA-dependent manner with SMN.

Secondly, we co-immunoprecipitated SMN with ribosomal proteins and translation factors from purified polysomes, and found that SMN is associated with ribosomal proteins through protein-protein interactions and with the Poly(A) binding protein (PABP) mainly through mRNA-dependent interactions (**Figure 1F**). This finding further suggests that SMN is preferentially associated to ribosomes/polysomes via mRNA-independent interactions.

Thirdly, to rule out the possibility that the observed interaction of SMN with ribosomes and polysomes derived from Gemin-granules (*32*, *33*), we compared the co-sedimentation profile of SMN, Gemin- and RNA-granules after sucrose gradient fractionation of cell lysates. We found that SMN co-sediments independently with a marker of Gemin-granules and with ribosomes, but not with mRNA-granules (**Supplementary Figure 2A**). Taken together, these findings suggest that, in addition to being part of Gemin-granules, SMN is a *bona fide* ribosome-associated factor.

Since SMN expression levels are known to be tissue-dependent (*34*), we wanted to establish whether SMN displays distinctive ribosome binding ability in a tissue-specific manner, dependent upon its relative abundance. To test this hypothesis, we established the relative co-sedimentation of SMN with ribonucleoparticles (RNPs), ribosomal subunits, ribosomes and polysomes in multiple tissues from wild-type mice: spinal cord, brain, kidney, liver and heart (**Figure 2A, B** and **Supplementary Figure 2B-D**). We observed that the association of SMN with the translation machinery is tissue dependent (**Figure 2C**). Interestingly, this variability is proportional with the overall abundance of SMN, whose level negatively correlates with RNPs association (r= −0.69) and positively correlates with 60S, 80S and polysome association (r = 0.78, 0.99 and 0.9 respectively) (**Figure 2D**). This confirms that the association of SMN with ribosomes and polysomes is tissue-specific and dependent on SMN concentration, as observed *in vitro* (**Figure 1B**). This finding suggests that a subset of ribosomes is associated with SMN in a concentration and tissue-dependent manner *in vivo*. We termed these ribosomes associated with SMN “SMN-primed ribosomes”.

**Figure 2.**
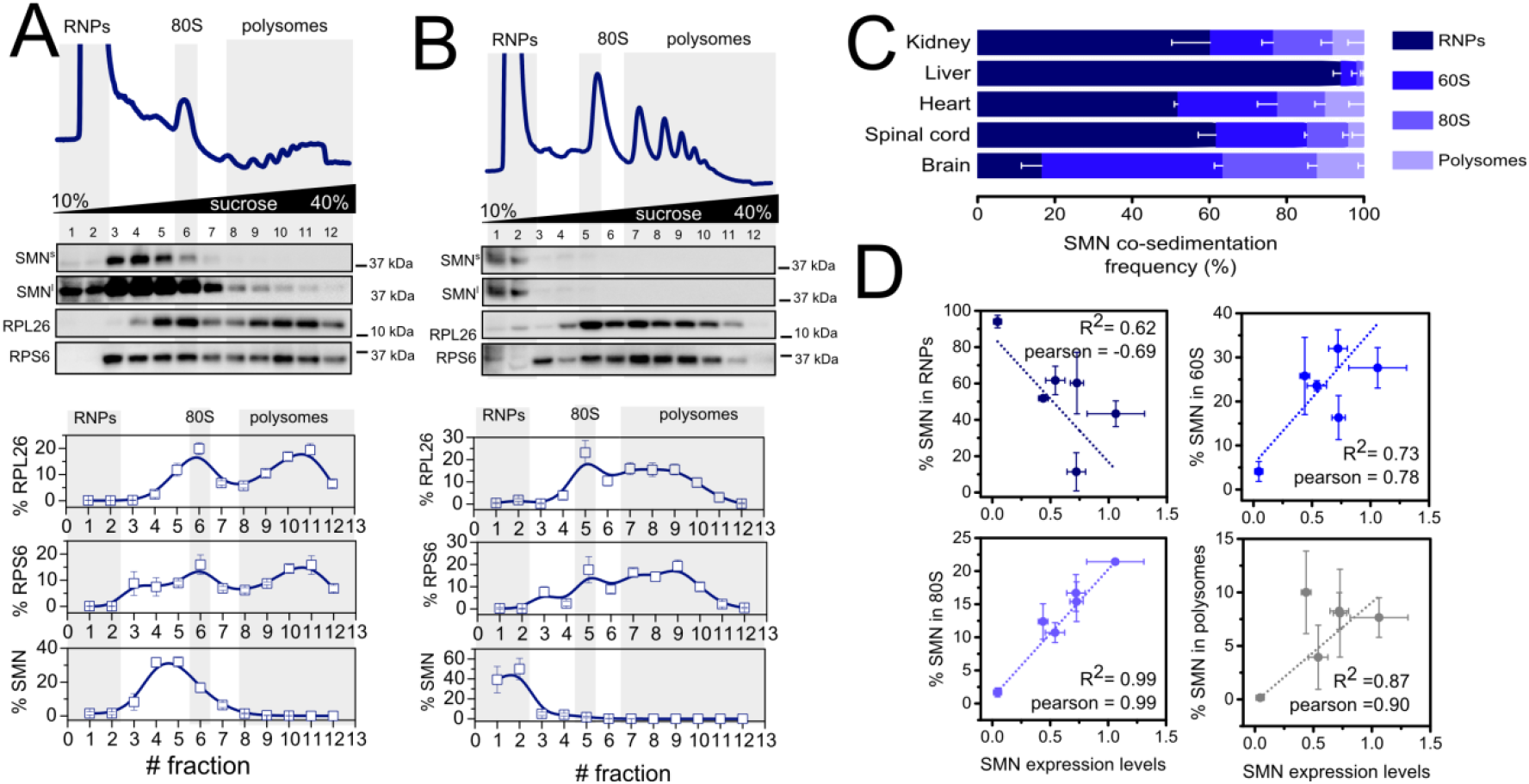
SMN interacts with RNPs and with the translation machinery in a concentration dependent manner across different tissues. Co-sedimentation profiles of SMN in brain (**A**) and (**B**) liver. The relative distributions of SMN, and markers of the small (RPS6) and large (RPL26) subunits of the ribosome were used as controls for sedimentation (upper panels). The relative distribution of each protein along the profile is shown as the average (± SE) of three independent replicates. (**C**) Summary of SMN co-sedimentation with RNPs, 60S, 80S and polysomes in different tissues. The percentages were obtained using co-sedimentation profiles shown in panels (**A**, **B** and **Supplementary Figure 2B-D**). All experiments were performed in biological triplicates. (**D**) Relationship between the relative expression level of SMN in different tissues obtained from Groen et al., 2018 (*34*) and the relative distribution of SMN in RNPs, 60S, 80S and polysomes obtained from (C).

### SMN positively regulates translation

It has previously been proposed that SMN may act as a repressor of cap-dependent translation *in vitro* (*28*), but this result is not in complete agreement with recent findings, including *in vivo* studies (*11*, *30*, *35*). Therefore, we wanted to establish whether SMN might be associated with stalled or actively translating ribosomes. To test this, we took advantage of an *in vitro* transcription translation assay using different concentrations of recombinant SMN and a reporter gene whose translation is controlled by an IRES sequence. We observed that a higher concentration of SMN leads to a higher production of the reporter protein (**Figure 3A**, **Supplementary Figure 3**), suggesting that SMN is a positive regulator of cap-independent translation *in vitro*.

**Figure 3.**
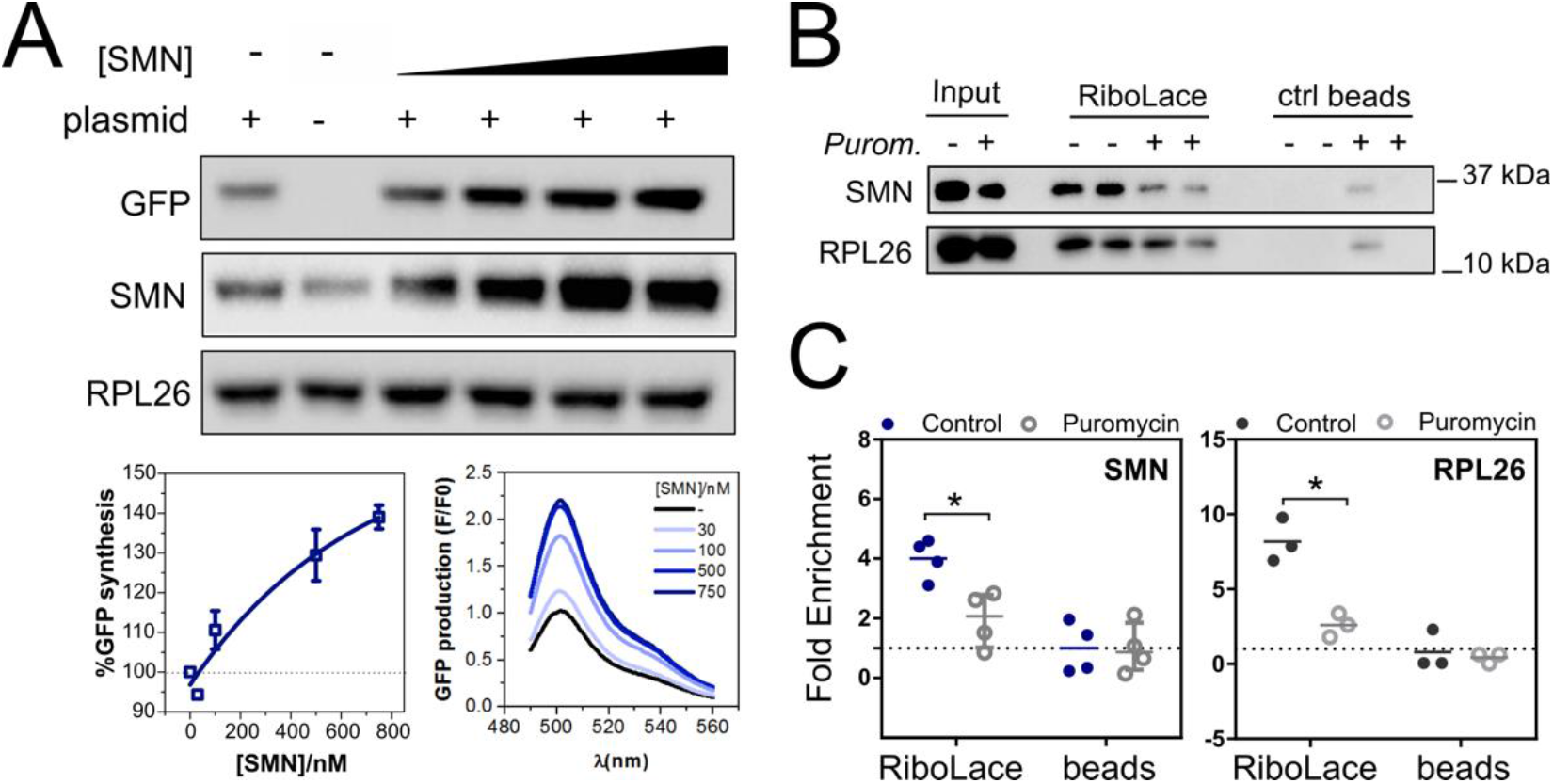
SMN is associated to actively translating ribosomes and positively regulates translation. (**A**) *In vitro* translation of reporter GFP in the presence of different concentrations of recombinant SMN. The abundance of GFP production was measured by western blotting to assess the impact of SMN addition on the *in vitro* translation systems. As a negative control a reaction in the absence of the GFP reporter was run in parallel. The ribosomal protein RPL26 was used as a loading control. Left lower panel, semi-quantitative analysis of GFP level in the presence of different concentrations of recombinant SMN. Plotted are the values ± s.d. from three independent experiments. Right lower panel, the production of GFP was also monitored by measuring the appearance of fluorescence in independent assays. (**B**) Western blot analysis of SMN association to active ribosomes using RiboLace (*36*) in human cells (upper panels, input, RiboLace and not-functionalized beads) before and after treatment with the translation inhibitor puromycin (100 μM, 1h). The ribosomal protein RPL26 is used as a marker of ribosomes. (**C**) The enrichment of SMN and RPL26 with respect to the not-functionalized beads is shown (n =3-4; t test; *p < 0.05).

Next, we checked the association of SMN to active ribosomes isolated *in cellulo* using the RiboLace method (*36*) (**Figure 3B** and **C**). We confirmed that SMN is associated with active ribosomes and released when translation is inhibited. These findings show that SMN positively regulates translation by binding to actively translating ribosomes.

### SMN-primed ribosomes are positioned within the first five codons of a specific subset of mRNAs

Having demonstrated that SMN binds to ribosomes and that SMN-primed ribosomes are connected to active translation, we next wanted to establish whether they control a specific subset of mRNAs and where SMN-primed ribosomes are preferentially positioned along transcripts. Therefore, we performed a ribosome profiling analysis of SMN-primed ribosomes in wild-type mouse brains (**Figure 4A**). We immunoprecipitated SMN from ribo-pellets exposed to RNase I. Notably, SMN co-immunoprecipitated with RPL26 (**Supplementary Figure 4A**), further demonstrating that SMN likely binds to ribosomes by RNA-independent interactions, as described above (**Figure 1E**). Next, we isolated and sequenced the RNA fragments protected by SMN-primed ribosomes, according to Ingolia et al., 2009 (*37*). In parallel, samples from immunoprecipitation with IgG were used as controls for possible unspecific isolation of ribosome protected fragments. By comparing RNA fragments protected by SMN-primed ribosomes to control IgG, we identified a set of 1095 transcripts, corresponding to 901 genes (**Supplementary File 1**). The vast majority of these transcripts (72%) are protein-coding (**Figure 4B**). Importantly, this experiment did not enrich for snRNAs or snoRNAs associated with the independent cytoplasmic role of SMN in RNP biogenesis, thus confirming that our analysis is specific for the subset of SMN bound to ribosomes.

**Figure 4.**
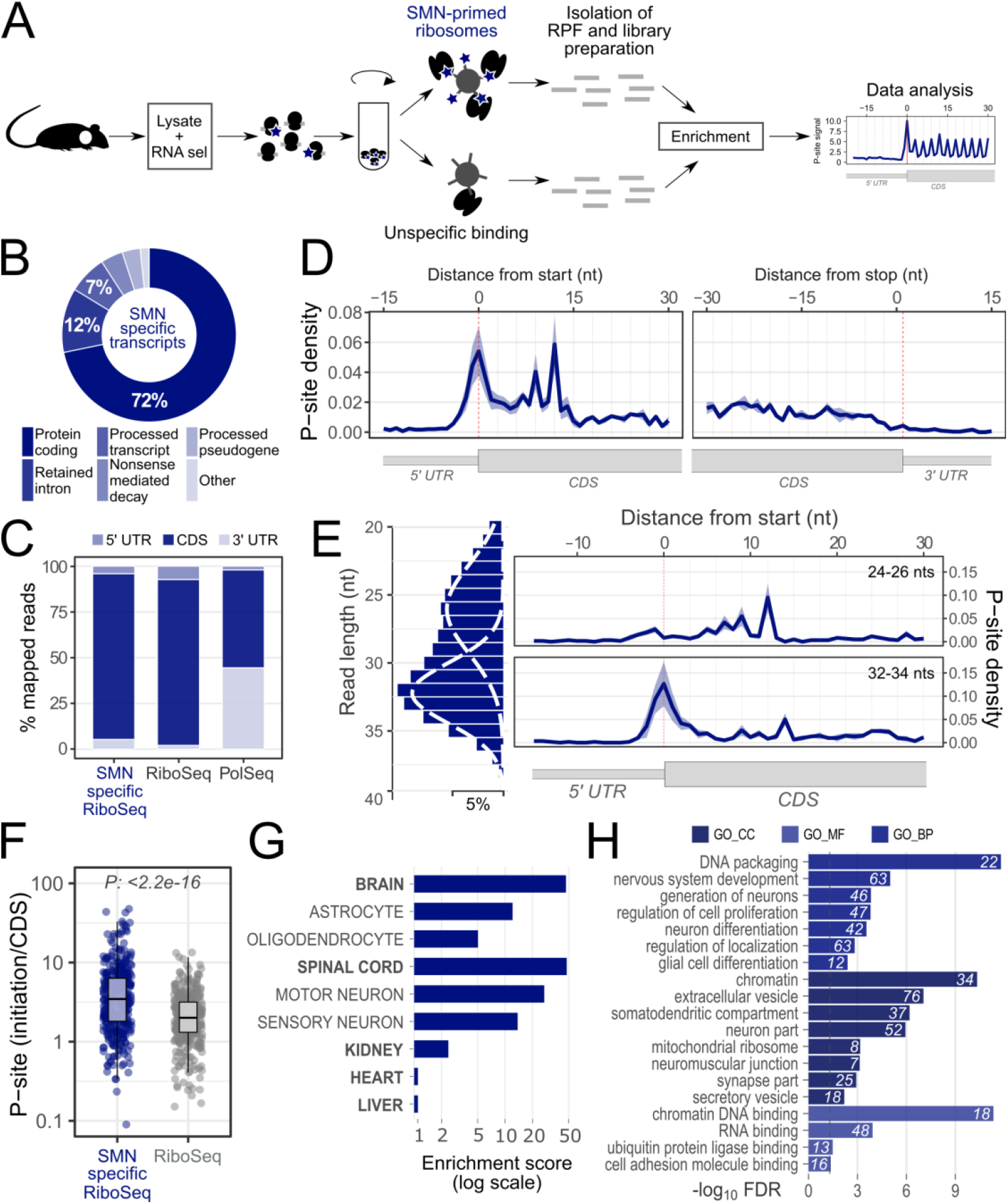
Ribosome profiling of SMN-primed ribosomes. (**A**) Schematic representation of the protocol used to isolate RNA fragments protected by ribosomes associated with SMN. In parallel, immunoprecipitation using IgG was performed to assess the unspecific capture of ribosome protected fragments. (**B**) Identification of transcript types enriched in fragments protected by SMN-primed ribosomes. SMN-primed ribosomes protect predominantly mRNAs. (**C**) Positional enrichment along the three mRNA regions of SMN-specific RiboSeq reads. The bar plots display the percentage of reads aligning on 5’ UTR, coding sequence and 3’ UTR for SMN-specific RiboSeq, RiboSeq and classical sequencing of polysomal transcripts (PolSeq from Bernabò et al., 2017 (*11*)) as control. All data were obtained from the same control brains. (**D**) Meta-profile based on the P-sites of reads mapping on mRNAs enriched in fragments protected by SMN-primed ribosomes. The graph depicts the average signal among replicates and the surrounding shadow represents the standard error. An accumulation of SMN-primed ribosomes on the first five codons of the CDSs emerges. (**E**) Two distinct populations of reads contribute to the accumulation of reads at the beginning of the CDS for mRNAs enriched in SMN-primed ribosomes. Left panel: distribution of read lengths, fitted with two Gaussian curves displayed as dashed lines. Right panels: meta-profiles based on the P-sites of short reads (24-26 nucleotides, upper panel) and long reads (32-34 nucleotides, lower panel). The lines report the average signals among replicates and the surrounding shadows represent the standard errors. (**F**) SMN-primed ribosomes preferentially occupy the beginning of the coding sequence of mRNAs. Dot plots showing the distributions of the ratios between the average number of P-sites on the first five codons (initiation) and the average number of P-sites on the whole coding sequence (CDS) for SMN-specific RiboSeq and classical RiboSeq of healthy mouse brains. The box plots associated with the distribution are also reported. Statistical significance was determined using a Wilcoxon-Mann-Whitney test (*** p-value < 2.2e-16). (**G**) Over-representation of tissue-specific markers among genes enriched in SMN-primed ribosomes. Tissues where SMN levels were previously measured are displayed. The enrichment score was calculated with enrichR. (**H**) Gene ontology and pathway enrichment analysis of genes enriched in SMN-primed ribosomes. The number of genes associated to the corresponding term is displayed on the right of the bars. (GO_BP:biological process, GO_CC:cellular component, GO_MF: molecular function).

To map the position of SMN-primed ribosomes, we identified the P-site location within SMN-primed ribosome protected fragments (RPFs) and calculated the percentage of P-sites falling within the coding sequence or UTRs of coding transcripts (**Figure 4C**). Remarkably, SMN-primed RPFs map prevalently to the coding sequence, similar to other ribosome profiling (RiboSeq) data and distinct from the sequencing of polysomal RNA (**Figure 4C**). This result demonstrates that SMN binds to ribosomes along the coding sequence, as expected for a ribosome profiling experiment.

Next, we analysed in detail the P-site position of SMN-primed RPFs near the start and stop codon of transcripts associated to SMN-primed ribosomes. We observed a clear accumulation of signal following the reading frame mostly within the first five codons of the coding sequence (**Figure 4D**). In particular, two distinct populations of ribosome protected fragments with different lengths contribute to this accumulation: a population of shorter fragments (24-26 nucleotides) peaking on the fifth codon (**Figure 4E**, upper panel), and a population of longer fragments (32-34 nucleotides) peaking on the first codon (**Figure 4E**, lower panel). These two populations are shared among the selected transcripts (**Supplementary Figure 4B**) and may be associated with different ribosome conformations, as previously suggested for yeast (*38*). We further confirmed that SMN-primed ribosomes preferentially occupy the beginning of the coding sequence by determining for each transcript the ratio between the number of P-sites on the first 5 codons (initiation) and the whole coding sequence (**Figure 4F**). Since the P-site signal is increasingly out of frame after the 5th codon (**Supplementary Figure 4C**), we considered the set of 385 protein coding transcripts displaying a signal within the first codons as *bona fide* SMN-specific transcripts for further analysis.

To gain insight into the possible role of these transcripts in different tissues, we analysed their tissue-expression patterns according to established datasets (*39*). Analysis of cell types within compartments of the nervous system reveals the highest enrichment for motor neurons, followed by sensory neurons, astrocytes and oligodendrocytes (**Figure 4G**). Outside the nervous system, we found a significant enrichment in kidney, but no enrichment in heart or liver (**Figure 4G**). Annotation enrichment analysis with Gene Ontology further highlights the significant association of neuron-specific functions with mRNAs enriched in SMN-primed ribosomes (**Figure 4H**).

### Transcripts bound by SMN-primed ribosomes display defects in ribosome recruitment at the third and fifth codons at early stages of SMA

To establish whether loss of SMN interaction with ribosomes gives rise to translational defects in SMA at early stage of disease (*11*), we first ruled out the possibility that translational changes are rather caused by pathways controlling translation, in particular the PERK (Unfolded Protein Response) and mTORC1 pathways. Both these pathways converge on polysomes, controlling the availability of initiation factors: eIF2alpha through the PERK pathway (*40*) and eIF4E through 4E-BP and the mTORC1 pathway (*41*). Since we did not observe any activation of these pathways at early stages of disease (**Supplementary Figure 5A**), the previously observed translational defects (*11*) (**Supplementary Figure 5B**) are caused by mechanisms independent of mTORC1 or PERK pathway inactivation/activation.

Next, we analysed the positioning of active ribosomes using the Active-RiboSeq method (*36*) and we compared SMA mouse brains to age matched controls (**Supplementary Figure 5C**). Notably, we observed that the majority of genes with altered translation (76%) are characterized by a decreased ribosome occupancy in SMA (**Figure 5A**). This strong translational down-regulation is in agreement with previously reported translation defects (*11*). Interestingly, whilst the 262 genes with increased ribosome occupancy in SMA do not display any tissue expression specificity, the 837 genes with defects in active translation are strongly enriched for both brain and spinal cord compartments, particularly in motor neurons (**Figure 5B**). Outside the nervous system, transcripts with decreased translation in SMA are significantly enriched in kidney, but not in heart or liver (**Figure 5B**). Importantly, this signature of tissue expression closely matches the one observed for SMN-specific transcripts (**Figure 4G**). Prompted by this finding, we further verified that SMN-specific transcripts display significantly decreased signal from Active-Riboseq in SMA, with respect to SMN-unspecific transcripts (P: 4.6e-11, **Figure 5C**).

**Figure 5.**
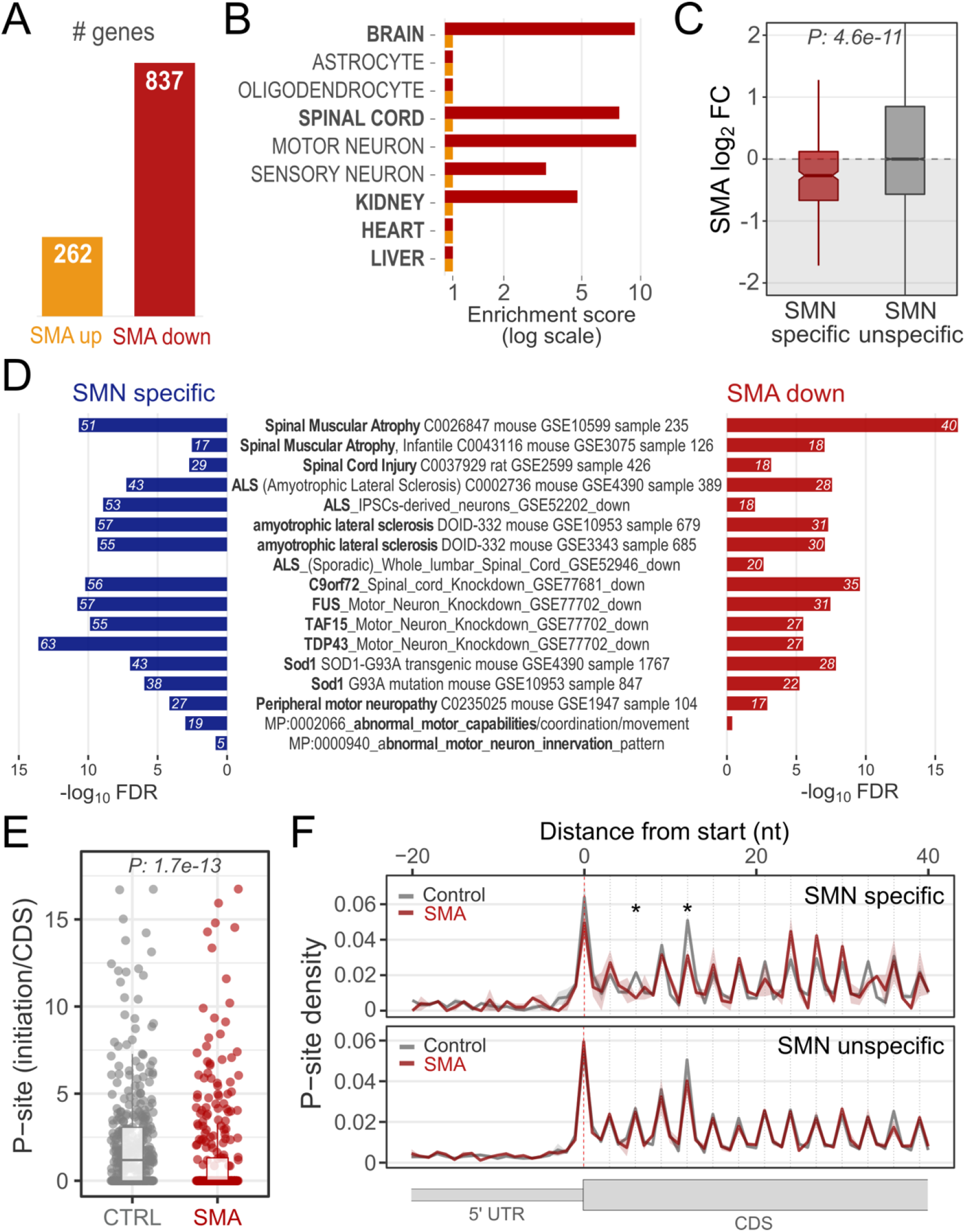
Transcripts bound by SMN-primed ribosomes display defects in positioning of active ribosomes at early stages of SMA. (**A**) Numbers of genes with significantly increased (up) or decreased (down) active ribosome occupancies in early symptomatic SMA mouse brain. (**B**) Over-representation of tissue-specific markers among genes with significantly increased (yellow) or decreased (red) active ribosome occupancies in SMA. Tissues where SMN levels were previously measured are displayed. The enrichment score was calculated with enrichR. (**C**) Comparison between SMA active ribosome occupancy changes in SMN-specific (red) and SMN unspecific (grey) genes. SMN specific genes show a significant shift towards a reduction in SMA active ribosome occupancy (Wilcoxon rank-sum test). (**D**) Over-representation analysis of terms associated with motor neuron diseases among genes enriched in SMN-primed ribosomes (SMN specific, blue) and genes with decreased translation occupancy in SMA (SMA down, red). Genes with increased translation occupancy in SMA did not show any significant enrichment for terms associated with motor neuron diseases. (**E**) Dot plots showing the distribution of the ratios between the average number of P-sites on the first five codons (initiation) and the average number of P-sites on the whole coding sequence (CDS) for SMN-specific transcripts based on control and SMA Active-RiboSeq signal. (**F**) Overlay meta-profiles based on the P-site position of active ribosomes comparing control (grey) and SMA (red) occupancies along SMN-specific transcripts (upper panel) and SMN unspecific transcripts (lower panel). Positional differences between control and SMA P-sites were tested using the T-test (* P-value < 0.05).

Importantly, transcripts bound by SMN-primed ribosomes and with defects in active translation in SMA match with transcripts previously associated with SMA and other motor-neuron diseases such as ALS (**Figure 5D**), suggesting that these experiments captured disease-relevant mRNA targets. In agreement with our previous finding that SMN-primed ribosomes are preferentially located around the first five codons (**Figure 4D-E**), SMN-specific transcripts exhibit lower active ribosome signals, specifically at the beginning of their coding sequence, as a result of SMN loss in SMA (P: 1.7e-13, **Figure 5E**). This is in keeping with a role for SMN in regulating the very initial phases of translation.

Given these observations, we wondered whether SMN-specific transcripts show positional defects in active ribosome P-sites within the first 5 codons. Comparing early symptomatic SMA and control tissues, SMN-unspecific transcripts do not display any significant difference in their active ribosome profiles (**Figure 5F** lower panel). Strikingly, however, SMN-specific transcripts are completely devoid of ribosomes at the 3rd codon and show a significant reduction in ribosome occupancy at the 5th codon (**Figure 5F** upper panel). These results suggest that loss of SMN induces positional defects in ribosome occupancy restricted to transcripts which are bound by SMN-primed ribosomes and that SMN is required during early stages of translation when the nascent chain is short and not yet deep inside the exit tunnel of the ribosome.

### Translationally defective transcripts in SMA display deficient use of rare codons in the coding sequence and enrichment in IRES in the 5’UTR

The association of SMN-primed ribosomes with the first five codons of a subset of mRNAs (**Figure 4D-F**), which are defectively translated in SMA (Figure 5C) and involved in SMA and ALS (**Figure 5D**), suggests a novel ribosome-based mechanism possibly underlying the pathophysiology and cell-type specificity of these diseases. Therefore, we explored in more detail the molecular and functional features characterizing these mRNAs. First, we found that they largely encode proteins either localized to the nucleus or mitochondria, including both transmembrane and secreted proteins (**Supplementary Figure 6A**). Then we found that these transcripts display shorter CDSs (median decrease of 194 nts, 19%), 5’ UTRs (decrease of 27 nts, 18%) and 3’ UTRs (decrease of 129 nts, 16%) (**Supplementary Figure 6B**). Consistently with the reported propensity of SMN to bind RNA sequences rich in G and A (*42*, *43*), GA-rich consensus motifs were found as enriched in both the 5’ and 3’UTRs (**Supplementary Figure 6C**) of SMN-specific transcripts. In addition, we found that the 5’ UTR of SMN-specific transcripts are enriched for IRES sequences (**Supplementary Figure 6D**), in agreement with our *in vitro* results. Consistently, transcripts with IRES sequences in their 5’ UTRs are more apt to show translational defects in SMA (**Figure 6A**).

To further verify that IRES sequences are involved in SMN-dependent translational control, we used a dual luciferase assay and followed the translational level of the reporter gene under the control of the well-known c-Myc IRES sequence (**Figure 6B**, upper panel). We tested the luciferase assay in motor neuron-like cell lines expressing SMN at either 100% or 20% of control levels (**Supplementary Figure 6E**). The 20% NSC-34 cell line recapitulates an *in cellulo* model of SMA, as it is characterized by a decrease in the fraction of ribosomes in polysomes (*11*) (**Supplementary Figure 6E**) similar to the one observed in SMA tissues (*11*) or after SMN silencing (*30*). Comparing NSC-34 cells with different SMN expression levels, with or without treatment with rapamycin (an inhibitor of cap-dependent translation), we found that cap-independent translation is impaired in a rapamycin independent manner when SMN expression is decreased (**Figure 6B**, lower panel), confirming a positive role of SMN in IRES dependent translation.

**Figure 6.**
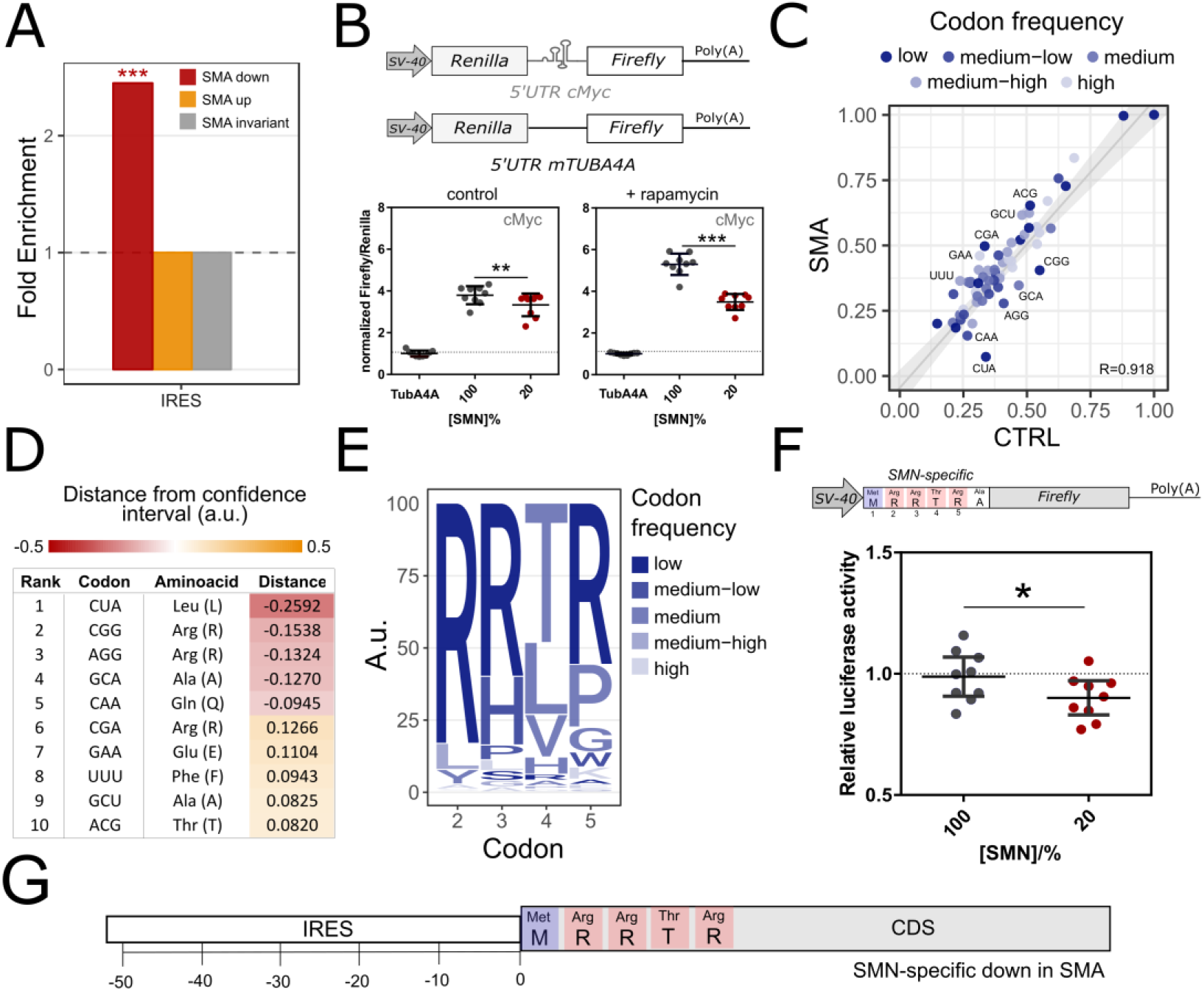
Translationally defective transcripts in SMA display deficient use of rare codons in the coding sequence and enrichment in IRES in the 5’UTR. (**A**) Over-representation analysis of IRES elements among genes with significantly increased (yellow) or decreased (red) active ribosome occupancies in SMA. IRES annotation was retrieved from Weingarten-Gabbay et al., 2016 (*71*). (**B**) SMN levels influence IRES-dependent translation revealed using a dual luciferase assay. Schematic representation of the pRuF-based bicistronic reporter vectors (top) containing Renilla luciferase cDNA (light gray) under the control of the constitutive promoter pSV40 and c-MYC 5′UTR IRES site placed upstream of the Firefly luciferase cDNA (white) (*65*). The 5’UTR of TubA4a was used as control. Luciferase assays were performed in NSC-34 cells lines (NSC-34 cells expressing 100% or 20% SMN) transiently transfected with the constructs. Results are shown as the ratio between Fluc and Rluc normalized to pRuF-Tuba4a vector, which is set to 1. Experiments were performed using three independent biological replicates and three technical replicates. Results are shown as the average (± SE); (n =3; t test; **p < 0.01; ***p < 0.001). (**C**) Comparison between the codon usage index, based on the sum of in-frame P-sites from control and SMA Active-RiboSeq. Each dot represents a codon and is colored according to the aminoacid frequency in the mouse transcriptome, divided in 5 classes (low: rare codons; high: frequent codons). The regression line, its 99% confidence level interval and the Pearson correlation coefficient are also displayed. Only the 10 furthest points from the confidence level are labelled. (**D**) Table reporting the triplet, the corresponding amino acid and the distance from the regression line of the codons. Negative values correspond to codons whose P-site coverage is lower in SMA than in the CTRL, positive values correspond to codons whose P-site coverage is higher in SMA than in the CTRL. (**E**) Logo-like representation of the most frequent amino acids codified by SMN-specific mRNAs with significant alterations in active ribosome occupancy in SMA at the beginning of the coding sequence. The enrichment analysis based on the number of occurrences of each codon was performed using brain-specific transcripts previously identified (see section Data Analysis of the manuscript) as background. Triplets with fold enrichments > 1 were selected and the weighted sum among synonymous codons was computed. The resulting values are displayed as percentages. Letters are colored according to the aminoacid frequency in the mouse transcriptome, divided in 5 classes (low: rare codons; high: frequent codons). (**F**) Luciferase assays for testing the contribution of first five codons features identified within the group of SMN-specific mRNAs exhibiting translational defects in SMA. Upper panel, schematic representation of the reporter firefly luciferase vector with the additional five codons. The reporter Renilla luciferase was used as transfection control. Luciferase assays were performed in NSC-34 cells lines (NSC-34 cells expressing 100% or 20% SMN) transiently transfected with both constructs. Results are shown as the ratio between Fluc and Rluc normalized to NSC-34 cells expressing 100%, which is set to 1. Results are shown as the average (± SE); (n =9; t test; **p < 0.01; ***p < 0.001). (**G**) Schematic diagram summarizing the combination of features in the 5’UTR and coding sequence of SMN-specific transcripts with translational defects in SMA.

In parallel, we analyzed the composition of the coding sequence of SMN-specific mRNAs and we surprisingly found that the first five codons are enriched in rare arginine codons with respect to SMN-unspecific mRNAs (**Supplementary Figure 6F**). Prompted by these findings, we further analysed the codon usage of mRNAs with translational impairment at the early stages of SMA (**Figure 5A**) and we observed a deficient usage of rare codons (**Figure 6C**), primarily encoding leucine and arginine amino acids (**Figure 6D**). Strikingly, when analyzing the codon usage of the first 5 codons in SMN-specific transcripts with significant translational defects in SMA, we found that arginine is the most frequent amino-acid (**Figure 6E**). This region of 5 codons matches precisely the position preferentially bound by SMN-primed ribosomes (**Figure 4D**) and shows a reduction of ribosome occupancy in SMA (**Figure 5E-F**). To confirm that this motif is sensitive to SMN loss, we used a luciferase assay (**Figure 6F**, upper panel) in motor neuron-like cells with different SMN expression levels. We found that the presence of the “arginine-rich” motif associated with SMN-specific transcripts induces a translational repression of the reporter in 20% SMN NSC-34 cells, validating the functional relevance of this sequence feature (**Figure 6F**, lower panel).

Taken together, these findings demonstrate that a combination of two features is required for SMN-specific mRNAs to be controlled translationally: i) IRES-like structures in the 5`UTR, and ii) a strong signature in the first five codons of the CDS, with rare codons, encoding arginine in particular, being the most frequent (**Figure 6G**).

### Acetylcholinesterase mRNA is an SMN-specific transcript and an early marker of local defects at the NMJ in SMA

To further explore the relevance of ribosome-based defects to SMA disease pathogenesis, we focused on transcripts bound by SMN-primed ribosomes and showing significant alterations in active ribosome occupancy in SMA (**Table 1**). We classified these targets according to their specific expression in tissues and cell types of the central nervous system (*39*, *44*) and found that, among the neuron specific genes, acetylcholinesterase (AChE) displays a significant decrease in translational efficiency not only at early stages of disease, but also at late stages in both brain and spinal cord (**Supplementary Figure 7A**). Importantly, in SMA mice treated with a single injection of ASO which restores SMN levels (*11*), AChE expression was restored to control levels, supporting a direct relationship between AChE expression and SMN levels (**Supplementary Figure 7A**).

**Table 1:** List resulting from the intersection of genes significantly enriched for SMN-primed ribosomes and genes with significant alterations in active ribosome occupancy in SMA. All genes displayed a decrease in active ribosome occupancy in SMA (P: 0.036, based on proportion test, P: 0.030 based on binomial test). Expression patterns were determined from Cahoy et al., 2008 (*44*) (CNS, neuron, astrocyte, ODC) and Lachmann et al., 2018 (*39*) (brain, spinal cord, motor neuron, sensory neuron).

| Table 1. List of genes significantly enriched for SMN-primed ribosomes with significant alterations in active ribosome occupancy in SMA. |  |  |  |  |  |  |  |  |  |  |
| --- | --- | --- | --- | --- | --- | --- | --- | --- | --- | --- |
| Gene | Description | SMA log2 FC | CNS | Neuron | Astrocyte | Oligodendrocyte | Brain | Spinal Cord | Motor neuron | Sensory neuron |
| Ache | acetylcholinesterase | -1.66 | P | P | NA | P | P | P | P | P |
| Lrrn1 | leucine rich repeat protein 1, neuronal | -1.59 | P | P | P | P | P | P | P | P |
| Chga | chromogranin A | -0.94 | P | P | P | P | P | P | P | P |
| Slc17a6 | solute carrier family 17, member 6 | -0.92 | P | P | P | NA | P | P | P | P |
| Peg3 | paternally expressed 3 | -0.93 | P | P | P | P | P | P | NA | NA |
| Acaa2 | acetyl-Coenzyme A acyltransferase 2 | -1.91 | P | P | P | P | NA | NA | NA | NA |
| D10Jhu81e | DNA segment, Chr 10 | -1.52 | P | P | P | P | NA | NA | NA | NA |
| Rcn2 | reticulocalbin 2 | -1.44 | P | P | P | P | NA | NA | NA | NA |
| Nxph4 | neurexophilin 4 | -1.72 | NA | NA | NA | NA | P | P | P | NA |
| Hist1h3i | histone cluster 1, H3i | -0.83 | P | NA | NA | P | NA | NA | NA | P |
| Hist1h2ah | histone cluster 1, H2ah | -1.00 | NA | NA | NA | NA | NA | NA | NA | NA |
| Hist1h3c | histone cluster 1, H3c | -0.94 | NA | NA | NA | NA | NA | NA | NA | NA |
| Get4 | golgi to ER traffic protein 4 homolog | -8.38 | NA | NA | NA | NA | NA | NA | NA | NA |
| Gm10094 | predicted gene 10094 | -1.47 | NA | NA | NA | NA | NA | NA | NA | NA |

P denotes the presence of expression determined according to Cahoy et al, 2008 (CNS, neuron, astrocyte, ODC) and Lachmann et al., 2018 (brain, spinal cord, motor neuron, sensory neuron). NA denotes the absence of expression.

By analyzing the sequence of the AChE transcript, we found that it recapitulates the sequence features characterizing SMN-specific mRNAs. In fact, the 5’ UTR of AChE is short and contains two putative short IRES-like sequences (**Figure 7A**). Additionally, the coding sequence of AChE codes for arginine in the second codon (**Figure 7A**) and other amino-acids encoded by rare codons which are also enriched in SMN-specific transcripts (**Figure 6E**). To confirm that these features are associated with translational defects under SMA-like conditions *in cellulo*, we performed two luciferase assays to evaluate the SMN-dependent translational control mediated by the 5’UTR region (**Figure 7B**) and by the sequence of the first five codons of the AChE CDS (**Figure 7C**), respectively. For both these functional motifs, we confirmed that loss of SMN expression causes translational defects in protein production of the reporter gene (**Figure 7B-C**).

**Figure 7.**
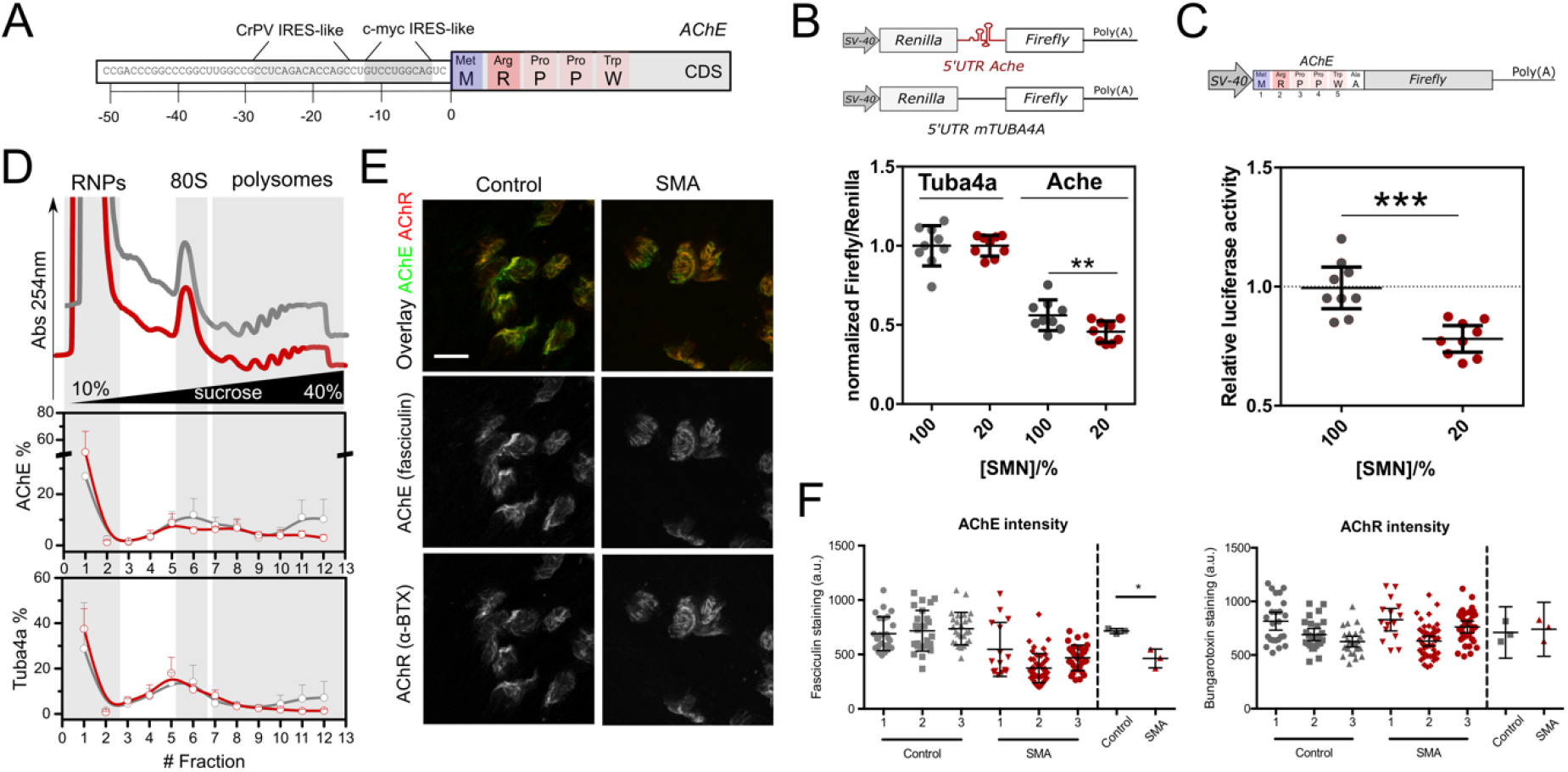
The acetylcholinesterase transcript shows ribosome drop-off and defective production of protein at the NMJ in SMA. (**A**) Schematic diagram summarizing the combination of features in the 5’UTR and coding sequence of AchE. The presence of IRES sequences in the AChE 5’UTR from CrPV and c-myc was revealed using IRESite (*72*). (**B**) SMN levels influence translation of a dual luciferase reporter bearing the 5’UTR of Ache. The 5’UTR of Tuba4a was used as control. Luciferase assays were performed in NSC34 cells lines (NSC34 cells expressing 100% or 20% SMN), transiently transfected with the constructs. Results are shown as the ratio between Fluc and Rluc (Relative Light Units, RLU) normalized to the results obtained with the pRuF-Tuba4a vector (where the empty value is set to 1). The experiments were performed in three independent biological replicates and three technical replicates. Results are shown as the average (± SE). Significant changes between cell lines was assessed using t-test (** p-value <0.01). (**C**) Luciferase assays for testing the contribution of the first five codons of AChE. Upper panel, schematic representation of the reporter firefly luciferase vector with the additional five codons. The experiment was performed as in Figure 6C Results are shown as the ratio between Fluc and Rluc normalized to NSC-34 cells expressing 100%, which is set to 1. Results are shown as the average (± SE); (n =9; t test; **p < 0.01; ***p < 0.001). (**D**) Relative co-sedimentation profile of Ache and Tuba4a along the sucrose gradient fractions of control (gray lines) and early symptomatic (red lines) brains. Data are represented as mean ± SE (n=3-4). (**E**) Representative confocal micrographs showing neuromuscular junctions in control (left panels) and late-symptomatic SMA mouse (right panels) FDB muscle. Acetylcholine receptors were labelled using alpha-bungarotoxin (BTX) conjugated to Alexa fluor 594 (red) and AChE was labelled using fasciculin-2 (FCC) conjugated to Alexa fluor 488 (green). Top panels show FCC / BTX overlap, middle and bottom panels show individual channels in greyscale. (**F**) FCC and BTX average intensity were determined for 2 FDB muscles in 3 control and 3 SMA mice. * *P* = 0.0295. AChR, acetylcholine receptor; AChE, acetylcholinesterase; a.u., arbitrary units. Scale bar: 10 μm.

According to these results and to our genome-wide data (**Figure 5D**), SMN-dependent translational defects are compatible with a loss of SMN-primed ribosomes along the coding sequence of SMN-specific transcripts. To test this further, we performed a co-sedimentation analysis of AChE mRNA after sucrose gradient fractionation in control and SMA tissues. Any change in the relative localization of the AChE transcript towards the lighter fractions (i.e. the RNPs) would indicate ribosome loss. Indeed, we observed that at an early stage of disease the AChE transcript is depleted from polysomal fractions, shifting towards the ribosome-free fractions (**Figure 7D**). In contrast, Tuba4a, which is not bound by SMN-primed ribosomes, does not show any change.

Finally, to confirm that the decrease in ribosomes along the AChE transcript leads to defective protein production during SMA *in vivo*, we observed that local expression of AChE protein is significantly impaired at the neuromuscular junction (NMJ) in symptomatic SMA mice (**Figure 7E-F**), whilst acetylcholine receptor (AChR) expression remains unchanged (**Figure 7E-F**). Downregulation of the AChE protein at the NMJ temporally follows the translational defect in ribosome occupancy at early stages (**Supplementary Figure 7B-C**), serving as a molecular marker of impairment at the NMJ in SMA.

## Discussion

Evidence accrued over recent years suggests that ribosomes are important players in fine-tuning translation, depending on their number (*45*), heterogeneity (*15*, *17*) and the binding of specific ribosome-associated factors (*19*, *46*). We explored the hypothesis that SMN is a ribosome-associated factor and demonstrated that SMN protein can be found in two distinct functional ‘populations’ within the cytoplasm. One population is associated with the well-known, canonical role of SMN in RNP biogenesis via its association with Gemin granules (*47*, *48*). According to Jablonka et al., 2001 (*49*) we found a second population which is not associated with Gemin granules, but rather associated with ribosomes, revealing SMN as a ribosome-associated factor *in vitro* and *in vivo*. This role is in agreement with SMN’s known ability to co-sediment with polysomes both *in vivo, in cellulo* and *in vitro* (*11*, *28–30*), as well as with general factors of translation such as eEF1A (*50*). Strikingly, the population of ribosomes associated with SMN, that we termed “SMN-primed” ribosomes, only made up a small fraction of the total ribosome pool. SMN-primed ribosomes display two unique characteristics: i) they are associated with a specific subset of mRNAs, and; ii) they are preferentially placed within the first five codons of the coding sequence.

SMN-primed ribosomes bind mRNAs characterized by strong enrichment in rare codons at the beginning of the coding sequence, particularly arginine-codons, and in IRES-like sequences in the 5’UTR. This ability to associate with a selected pool of mRNAs has been previously observed for ribosomes containing particular ribosomal proteins (*15*, *20*). Notably, we found that a significant number of mRNAs bound by SMN-primed ribosomes have been previously linked to the pathogenesis of SMA, as well as to related neuromuscular conditions such as ALS. This provides a molecular explanation as to why defects in ubiquitously-expressed proteins, such as SMN, can lead to a targeted degeneration of motor neurons, as observed in both SMA and ALS.

The fact that SMN-primed ribosomes are located within the first codons of SMN-specific transcripts suggests a highly specific, local function for this defined subpopulation of ribosomes. A general enrichment of signal at the beginning of the coding sequence is a property shared with ribosomes in general, as previously observed in several studies using classical ribosome profiling (*51–53*). Nonetheless, in the case of SMN-primed ribosomes, the signal is almost exclusively located at the very beginning (i.e. within the first 5 codons), and SMN-specific transcripts show a definite enrichment in rare codons in those positions. Importantly, it has been observed that a ribosome-pause at these very first codons acts as a translational-checkpoint to ensure productive ribosome elongation and protein synthesis (*53*). Accordingly, SMN binds to the elongation factor eEF1A (*50*) and we found that it is required for productive translation *in vitro*. This positive regulation of translation is lost *in vivo* and *in cellulo* when SMN expression is reduced (*11*, *28*, *30*). In addition, analyzing the read lengths of mRNA fragments protected by SMN-primed ribosomes, we found two distinct populations of reads characterized by different lengths. Similar bimodal distributions in read lengths have been observed in yeast, and have been associated with diverse structural conformations of the ribosome during the elongation phase of translation (*38*, *54*), consistent with our current observations. Thus, we propose a model whereby SMN regulates the translation of rare codons by acting as a stabilizer of specific ribosome conformations at the very beginning of the coding sequence, where it can induce a functional ribosome slowdown that ensures productive translation (**Supplementary Figure 8A**).

By profiling actively translating ribosomes (*36*) in control and early symptomatic SMA tissues we found that the vast majority of genes associated with a significant variation in ribosome occupancy displays a strong decrease in the number of active ribosomes upon SMN depletion, in agreement with previous studies (*11*). Strikingly, the mRNAs associated with SMN-primed ribosomes showed profound positional defects at the third and fifth codon, compatible with a loss of ribosome pausing mediated by SMN-primed ribosomes at a critical initial step during translation as in Han et al., 2014 (*53*). Importantly, rare codons show significantly reduced ribosome occupancy in SMA, with arginine-codons being once again the most affected.

Ribosome drop-off has been shown to occur with higher probability in the absence of a translational pause around the 5th codon (*53*). Consistently, loss of SMN-primed ribosomes at the fifth codon in SMA is associated with an overall decrease in ribosome occupancy at rare codon positions, which is suggestive of specific defective ribosome recruitment and ribosome drop off. These results lead us to propose a mechanistic model explaining how translational defects occur in SMA (**Supplementary Figure 8A**).

Among the mRNAs bound by SMN-primed ribosomes and characterized by translational defects in SMA, we identified acetylcholinesterase (AChE) which embodies most of the sequence features we identified in SMN-specific transcripts. AChE performs a central role in NMJ function by turning over acetylcholine after it has been used for signal transduction by motor neurons. Dysfunction and denervation of the neuromuscular junction (NMJ) is one of the earliest pathological features of SMA (*55–57*). Although NMJ pathology has been well-characterized at the cellular level, the molecular mechanisms associated with NMJ pathology remain poorly understood. In agreement with our proposed model, we found that, in early symptomatic SMA, the recruitment of AChE mRNA on polysomes is reduced and that this decrease is followed by defects at the protein level in the neuromuscular junction at later stages of disease. Previously, an absence of the asymmetric A12 form of AChE was observed in the serum of SMA Type I patients (*58*). Moreover, mutations affecting AChE in humans cause congenital endplate acetylcholinesterase deficiency. This disease displays a number of clinical features overlapping with those observed in SMA, from the age of onset and spinal deformity to muscle weakness and respiratory distress (*59*), suggesting possible common underlying pathological mechanisms. Thus, SMN-primed ribosomes play a crucial role in regulating AChE levels likely to contribute significantly to NMJ defects that are core to the pathogenesis of SMA.

The robust influence of SMN levels on ribosome binding, alongside the higher relative concentration of SMN protein found in the nervous system (*34*), supports a model whereby a stronger effect on transcripts bound by SMN-primed ribosomes should be observed in these tissues. Thus, a novel scenario for better understanding the molecular pathogenesis of SMA can be generated (**Supplementary Figure 8B**), in which tissue- and concentration-specific regulation of SMN concertedly tune the correct translation of mRNAs bound by SMN-primed ribosomes, as illustrated here by the effect observed on AChE.

Taken together, our findings demonstrate a central role for SMN in the regulation of ribosome heterogeneity, acting as a master modulator of ribosome fluxes on a disease-specific subset of disease-relevant mRNAs characterized by specific sequence features. This reveals an important role for ribosome-associated factors in the regulation of tissue-specific disease pathogenesis in SMA and related conditions.

## Materials and Methods

### Animal models

All animal procedures and breeding were performed in accordance with University of Edinburgh institutional guidelines and under appropriate project and personal licenses granted by the UK Home Office (PPL: P92BB9F93). The ‘Taiwanese’ mouse model of severe SMA (*60*), on a congenic FVB background, was established from breeding pairs originally purchased from Jackson Labs. ‘Taiwanese’ SMA mice (*Smn*−/−; *SMN2tg/0*) carry two copies of *SMN2* on one allele on a null murine *Smn* background. Phenotypically normal heterozygous (*Smn*−/+;*SMN2tg/0*) littermates were used as controls. Mice were maintained following an established breeding strategy (*61*). Litters were retrospectively genotyped using standard PCR protocols. All mice were housed within the animal care facilities in Edinburgh under standard SPF conditions. For clarity, throughout the paper we refer to the time points at which tissue was collected as early and late symptomatic. Early symptomatic was postnatal day 5 (P5) and late symptomatic was P7. All tissues were quickly dissected, snap-frozen and stored at −80°C until use.

### Subcellular fractionation from tissues

The method was adapted from Francisco-Velilla et al., 2016 (*31*). Briefly, half brain or whole spinal cord preparations were pulverized under liquid nitrogen with mortar and a cytoplasmic lysate was obtained according to Bernabò et al., 2017 (*11*). A few μL of the lysate was kept for protein extraction, representing the input fraction. The remaining sample was centrifuged for 67 min at 100,000 rpm using a TLA100.2 rotor (Beckman). The supernatant corresponds to proteins not associated to ribosomes or polysomes (unbound). The pellet containing ribosomes, (R pellet) was solubilized in 15 mM Tris-HCl pH 7.4, 100 mM KCl, 5 mM MgCl2, 2 mM DTT, 290 mM sucrose. A few μL were kept for protein extraction, the remaining volume was adjusted to KCl 500 mM, to dissociate proteins mildly associated with ribosomes. The sample was subsequently loaded on a discontinuous sucrose gradient (720 μL buffer 40% (w/v) sucrose, 15 mM Tris-HCl pH 7.4, 500 mM KCl, 5 mM MgCl2, 2 mM DTT (bottom layer) and 480 μl buffer 20% (w/v) sucrose, 15 mM Tris-HCl pH 7.4, 500 mM KCl, 5 mM MgCl2, 2 mM DTT (top layer) and ultra-centrifuged at 100,000 rpm or 1.5 hrs using a TLA100.2 rotor (Beckman). The supernatant contains proteins loosely associated to ribosomes and polysomes (LBR), while the pellet contains washed ribosomes (RBR). The RBR pellet was dissolved in sample buffer for SDS-PAGE electrophoresis and western blotting. The proteins in the other fractions were extracted using a Methanol/Chloroform protocol. In the case of RNAse treatment, the R-pellet prepared as described above was treated with RNAse I (1.5U/1Abs260) for 45 min at room temperature. After RNAse treatment, the lysate was ultracentrifuged as described above to isolate the proteins associated to ribosomes/polysomes via RNA dependent interactions (supernatant, RNA-dep) and proteins associated via protein-dependent interactions (pellet, protein-dep). Protein extraction was performed as described above before analysis by SDS-PAGE electrophoresis or western blotting.

### Western blotting from tissues

Protein expression in tissue from Taiwanese SMA mice was performed using standardized protocols as described previously (*62*). Briefly, tissues were homogenized on ice in RIPA buffer with 1x protease inhibitor. Protein concentration was normalized using a BCA assay. Proteins were size-separated on a 4-12% Bis-Tris gradient gel and transferred to PDVF membrane using an iBlot semi-dry blotting system. Membranes were blocked in blocking buffer (LI-COR) and incubated overnight in primary antibodies. Antibody detection was performed using fluorescent secondary antibodies (LI-COR) and protein loading was normalized to a fluorescent total protein stain (LI-COR). Quantification was performed as described previously (*34*).

### Ribosome purification and ribosome-binding assay

Purification of 80S ribosomes was performed from NSC-34 cells depleted by SMN using CRISPR-Cas9 technology. A clone of NSC-34 cells with 0% SMN expression level was used to obtain a cytoplasmic lysate as described previously (*11*). To enrich the sample for 80S ribosomes, the lysate was treated with RNase I (1.5 Units /1 Abs260 lysate) at RT for 45 min and analysed by sucrose gradient (10-40%) according to Viero et al., 2015 (*63*). Fractions corresponding to the 80S peak were collected. To purify ribosomes, 2mM DTT was added to the 80S sucrose fraction which was then centrifuged at 90,000 rpm for 4 h using a TLA100.2 rotor (Beckmann). The 80S pellet was resuspended in 10 mM Tris–HCl pH 7.5, 10 mM MgCl2, 150 mM NaCl, 2 mM DTT, 100 ug/mL cycloheximide and stored at −80°C. Ribosome concentration was calculated as Abs260 = 1 A.U. corresponds to [80S]= 20 nM (*64*). Recombinant SMN was purchased (ENZO) and incubated at varying ribosome:SMN molar ratios for 2 h at 4°C. SMN bound ribosomes were purified from unbound SMN by ultracentrifugation at 90,000 rpm for 4 h using the TLA100.2 rotor (Beckmann). The supernatant was kept for protein purification by using the chloroform/methanol protocol and the pellet was directly dissolved in sample buffer (Santa Cruz), heated at 99°C for 10 min and resolved by SDS-PAGE. SMN and ribosomal markers (RPL26 and RPS6) were detected by WB using anti-SMN, anti-RSP6 (40S, 80S) or RPL26 (60S and 80S) antibodies. Independent binding assays were conducted at least three times.

### In vitro transcription-translation assays

*In vitro* transcription-translation was performed according to manufacturer’s instructions using the 1-Step Human Coupled IVT Kit HeLa lysates and pCFE-GFP as reporter (Thermo Scientific). The reporter contains a T7 promoter and an EMCV internal ribosome entry site (IRES) for cap-independent translation. For quantifying the protein synthesis levels, we adopted two approaches: western blotting and fluorescence spectroscopy. For western blotting, after incubation of each reaction at 30°C for 1.5 h, proteins were extracted by chloroform/methanol. The proteins were solubilized in Sample Buffer and analyzed by SDS-PAGE. The production of EGFP was monitored using an anti-TurboGFP Antibody (Thermo Scientific). The ribosomal proteins RPL26 and RPS6 were used as controls for ribosomes. Protein production was quantified by densitometric analysis using ImageJ. The EGFP signal was normalized to the RPL26 signal. To estimate the protein production by fluorescence spectroscopy, we measured the GFP amount from the height of the emission spectra maximum at 502 nm. 10 μL sample after 1.5 h incubation at 30°C in the presence of different SMN concentrations, prepared as described above, was added to a 1-cm quartz cuvette filled with 990 μL of buffer. Spectra were acquired on a Fluoromax-4 (Horiba Jobin-Yvon) with l_ex_=482 nm. Alternatively, samples were prepared in a 386-wells black microplate (Dynex), and fluorescence measurements immediately started at 30°C with FLUOstar Galaxy (BMG, Ortenberg, Germany), equipped with excitation and emission fluorescence filters centered at 480 and 510 nm, respectively. A reading point every 36’ for has been collected for the entire experiment duration (300min).

### Immunoprecipitation of SMN-associated proteins from polysomal fractions and ribosomal pellets

IP from polysomal fractions was performed on pooled polysomal fractions from control brains according to Bernabò et al., 2017 (*11*). To decrease the viscosity of the solution and increase the efficiency of the immunoprecipitation the samples were diluted 3x using 30 mM Tris-HCl (pH 7.5), 100 mM NaCl, 10 mM MgCl2, 20 μL/mL cycloheximide. One mL of pooled fractions was kept as input, and the remaining sample was divided into two parts and incubated for 2 h at 4 °C with either 2 μg of SMN antibody (BD Biosciences) or, as negative control, 2 μg of anti-Mouse IgG (Life Technologies) while slowly rotating. After the addition of a mix of Dynabeads Protein G and Dynabeads Protein A (Life Technologies), the samples were kept rotating for an additional 2h at 4 °C. After separation of the beads from the supernatant, which was stored as “Unbound”, the beads were washed three times with 500 μL Washing Buffer (10 mM Tris–HCl pH 7.5, 10 mM MgCl2, 10 mM NaCl, 1% Triton X-100, 1 mM DTT, 0.2 mg/mL cycloheximide, 1% Na-deoxycholate, 2.5 μL/mL Protease Inhibitor Cocktail). RNase A/T1 was the added (200 μg/mL final concentration) and the samples were kept under rotation for 1.5 h at 4 °C to separate molecules interacting with SMN via protein-protein interactions from molecules released by the nuclease. Samples were placed again on the magnetic stand and the supernatant was stored and proteins (“RNA-mediated”) were extracted. After extensive washing, the molecules attached to the beads (“Protein-mediated”) were directly dissolved in electrophoresis sample buffer for western blotting analysis. All other samples (“Input”, “RNA mediated” and “Unbound”) were purified by methanol/chloroform extraction and analyzed by western blotting with the protein-protein interactors.

### Cloning of 5’UTRs and SMN-specific motives and Luciferase assay

The first five codons of SMN-specific CDS and AchE CDS were cloned into a pGL3 luciferase reporter vector. Both motives were created by annealing complementary oligos forming overhangs of endonuclease restriction sites at the 5′- and 3’-ends (HindIII and NcoI, respectively; see **Supplementary Table S1**). The motives were inserted upstream of the *Firefly* luciferase start codon.

Cloning of the 5’UTRs of AChE 5’-UTR and TUBA4A 5′-UTR was performed into a bicistronic vector. Sequences were PCR amplified from cDNA obtained from mouse brain (P5) using primers that contained endonuclease restriction sites at their 5′- and 3’-end (EcoRI for Fw primer and NdeI for Rev primer; see **Supplementary Table S1**). The amplicon was then cloned into pRuF plasmid, downstream of the *Renilla* stop codon and upstream of the *Firefly* start codon. In the pRuF vector Renilla and Firefly cDNAs are transcribed as part of a single transcript separated by the 5′-UTR sequence. All plasmid clones were checked by DNA sequencing.

For luciferase assay 1 × 10^5^ NSC-34 native and NSC-34 expressing 20% SMN cells (*11*) were seeded in 24-well plates. In the case of pGL3 plasmids, cells were co-transfected in 1:3 ration with pRLSV40 plasmid to normalize for transfection efficiency. Luciferase assays were run after 24h. In the case of pRuF plasmids, which contain both Renilla and Firefly luciferases, luciferase assays were run 24-48h after transfection. Lciferase assays were performed using the dual-luciferase reagent (Promega) as previously described (*65*). For each cell line 4-9 independent biological replicas were analyzed.

### Immunoprecipitation of SMN with active ribosomes

Control MCF7 cells and MCF cells treated with puromycin 100 μM final concentration for 1h were lysed according to Clamer et al., 2018 (*36*). Active ribosomes were isolated using the RiboLace kit (Immagina Biotechnology) and proteins were extracted directly from beads using sample buffer (Santa Cruz). Five μL of lysate was kept as input. Proteins were resolved using SDS-PAGE and western blotting as in Clamer et al., 2018 (*36*).

### Active and classical Ribosome profiling

Cytoplasmic lysates were prepared by pulverizing frozen P5 control and early symptomatic SMA mouse brains, according to Bernabò et al., 2017 (*11*). To capture RNA fragments protected by active ribosomes (*36*), we used the RiboLace technology (Immagina Biotech) following manufacturer’s instructions. The indexes used for preparation of the library preparation of both classical and active ribosome profiling are listed in **Supplementary Table S2**. The libraries were purified from primer dimers and adaptor dimers on an 8% native polyacrylamide gel and their quality and quantity were assessed by using the high-sensitivity DNA chip on the BioAnalyzer (Agilent) according to the manufacturer’s protocol and Qubit® 2.0 (ThermoFisher Scientific). The libraries were sequenced on an Illumina HiSeq2500 by the Core Facility, NGS (University of Trento).

Classical ribosome profiling was performed after polysomal purification as follows. The tissue lysates from control and early symptomatic brain were loaded on a 15-50% sucrose gradients. Polysomes were purified by ultracentrifugation and collected using a Teledyne ISCO model 160 equipped with a UA-6 UV/ VIS detector. Polysomal fractions were pooled and digested with Rnase I (150U/unit of area of polysomes, calculated from the polysomes profile) for 2h at 4°C. The digestion was then stopped by adding 400 U SUPERase-In RNase inhibitor (Thermo Fisher Scientific). The RNA was extracted as in Bernabò et al., 2017(*11*). Ribosome Protected Fragments (RPF) were isolated and the libraries were prepared following the Ingolia protocol (*66*). Active ribosome profiling was performed in parallel to classical ribosome profiling, starting from 25 μL of tissue lysates treated with RNAseI (5U/ absorbance at 260 nm). The nuclease digestion was performed on the same brain lysates used for Ribo-Seq. Two biological replicates were performed.

### Ribosome profiling of SMN-primed ribosomes

P5 brains were pulverized under liquid nitrogen and half of each sample was dissolved in lysis buffer (10 mM Tris–HCl pH 7.5, 10 mM MgCl2, 10 mM NaCl, 1% Triton X-100, 5 U/mL DNase I, 600 U/mL RiboLock RNase Inhibitor (Thermo Scientific), 0.2 mg/mL cycloheximide, 1% Na-deoxycholate, Protease Inhibitor Cocktail). After centrifugation to remove tissue debris, nuclei and mitochondria, the absorbance at 260 nm was measured. The concentration of NaCl was adjusted to 150 mM and endonuclease digestion was performed with RNAse I (5U/Unit of absorbance at 260nm in the lysate) at room temperature for 45 min. The reaction was stopped with SUPERase-In RNase inhibitor (Thermo Fisher Scientific). The digested lysates were centrifuged for 70 min at 100,000 rpm (TLA100.2 rotor, 4°C) and the ribosome pellet solubilized in 10 mM Tris–HCl pH 7.5, 10 mM MgCl2, 150 mM NaCl, 1% Triton X-100, 600 U/mL RiboLock RNase Inhibitor (Thermo Scientific), 0.2 mg/mL cycloheximide, Protease Inhibitor Cocktail. Ribosomes associated to SMN were obtained by immunoprecipitation using mouse anti-SMN antibody or anti-IgG as a control for unspecific binding. Briefly, the ribosome suspension was incubated with 2 μL of antibody for 1 h and 40 min in orbital rotation at 4°C. Then 100 μL of Dynabeads Protein G (Life technologies) were added and the sample was incubated for 1 hr at 4°C in orbital rotation. The supernatant was removed using a magnetic stand and the beads were washed 2 times for 5 min with washing buffer (10 mM Tris–HCl pH 7.5, 10 mM MgCl2, 150 mM NaCl, 1% Triton X-100, 0.2 mg/mL cycloheximide, Protease Inhibitor Cocktail) before extraction of RNA with Trizol. The ribosome protected fragments from both SMN and IgG immunoprecipitation were isolated and used for library preparation as described above. Experiments were performed in triplicate.

### Data Analysis

Reads were processed by removing 5ʼ adapters, discarding reads shorter than 20 nucleotides and trimming the first nucleotide of the remaining ones (using Trimmomatic v0.36). Reads mapping on the collection of *M. musculus* rRNAs (from the SILVA rRNA database, release 119) and tRNAs (from the Genomic tRNA database: gtrnadb.ucsc.edu/) were removed. Remaining reads were mapped on the mouse transcriptome (using the Gencode M6 transcript annotations), antisense and duplicate reads were removed. All alignments were performed with Bowtie2 (v2.2.6) employing default settings.

Normalization among replicates was performed with the trimmed mean of M-values normalization method (TMM) implemented in the edgeR Bioconductor package. Transcripts used for further analyses were selected using a threshold on their signal (FPKM and CPM values > 80th percentile). Classical RiboSeq and Active-RiboSeq data were processed separately, due to the different protocols used for capturing mRNA ribosome protected fragments. A common population of brain-specific transcripts was identified by merging the mRNAs identified for each technique.

Ribosome profiling analyses based on positional information were performed using riboWaltz (*67*). Fold-enrichment analyses were performed with edgeR and SMN specific transcripts were selected with the following thresholds: fold enrichment > 2 with respect to IgG, P value < 0.05, normalized number of ribosome-protected fragment > 1 per million. Genes with significant alterations in active translation were selected with the following thresholds: absolute log2 fold change > 0.75, P value < 0.05, normalized number of ribosome-protected fragment > 1 per million.

Functional annotation enrichment analyses were performed with Enrichr (http://amp.pharm.mssm.edu/Enrichr/).

### Data availability

Raw and analyzed classical and active ribosome profiling data of SMA mouse brains have been deposited under GEO: XXX (RiboSeq), XXX (Active-RiboSeq). Ribosome profiling data from SMN-primed ribosomes of healthy mouse brains have been deposited under GEO: XXX.

Classical and active ribosome profiling data of healthy mouse brains were retrieved from GEO: GSE102318 (RiboSeq), GSE102354 (Active-RiboSeq).

### Co-sedimentation profiles of proteins and of mRNA by droplet-PCR

Polysomal profiling from cell lines or from tissues was performed according to previously described protocols (*11*). Cytoplasmic lysates were loaded on a linear 10%–40% [w/v] sucrose gradient and centrifuged in a SW41Ti rotor (Beckman) for 1 h 30 min using a SW41 rotor at 40,000 rpm in a Beckman Optima XPN-100 Ultracentrifuge. Fractions of 1 mL were then collected monitoring the absorbance at 254 nm with a UA-6 UV/VIS detector (Teledyne Isco). Proteins from each sucrose fraction were extracted and analysed by SDS-PAGE and western blotting as described in Bernabò et al., 2017 (*11*) and Tebaldi et al., 2018 (*68*).

RNA from each sucrose fraction was extracted using TRIZOL. Equal volumes of RNA were used for cDNA synthesis using iScript cDNA synthesis kit. EvaGreen-based ddPCR reaction mixtures (23 μL final volume) were composed of 1x QX200 EvaGreen ddPCR Supermix, 150 nM forward and reverse primers (**Supplementary Table 3**) and 10 μL of 1:10 diluted cDNA. A 20 μL aliquot from each of the assembled ddPCR mixtures and 70 μL Droplet Generation Oil for EvaGreen were loaded into an eight-channel disposable droplet generation cartridge (Bio-Rad). After sample partitioning with QX200 Droplet Generator (Bio-Rad), the entire droplet emulsion volume was transferred with a Rainin multichannel pipette into a 96-well plate, heat-sealed with a pierceable foil in the PX1 PCR plate Sealer (Bio-Rad), and placed in a T100 thermal cycler (Bio-Rad). Thermal cycling conditions included the following steps, all at 2 °C/sec ramp rate: 5 min at 95 °C, 40 cycles of 30 sec at 95 °C and 60 sec at 60 °C, 5 min at 4 °C, 5 min at 95 °C and finally 12 °C indefinite hold. After PCR, droplets were read individually by QX200 Droplet Reader (Bio-Rad) and the data analysed by QuantaSoft software (Bio-Rad). The % of each transcript distribution along the profile was obtained using the following formula:

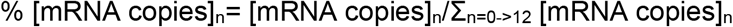

where n is the number of the fraction.

### Translation efficiency by qPCR

Total and polysomal RNA from control and early symptomatic brain, from late symptomatic brain and spinal cords (*11*) and from late symptomatic brain treated with ASO were purified as in Bernabò et al., 2017 (*11*). The retrotranscription reaction was performed starting from 100-500 ng of RNA using RevertAid First Strand cDNA synthesis kit (Thermo Scientific). qPCR was run on CFX Connect Real-Time PCR Detection System (BioRad) using Kapa Syber Fast qPCR Mastermix (Kapa Biosystems). Primer sequences are provided in **Supplementary Table 3**. Tuba4a was used as reference gene. All reactions were performed as two to three biological replicates and two to six technical replicates. The variation of translation efficiency (TE) was calculated as the difference between the fold change at the polysomal level and the fold change at the total level of the gene of interest in control versus SMA samples.

### NMJ fluorescence microscopy

For NMJ analysis, flexor digitorum brevis (FDB) muscle was dissected from early- and late-symptomatic mice using procedures similar to those described previously (*69*) and fixed in 4% PFA for 20 min at RT. After fixing, muscles were cleaned of remaining connective tissue and tendons. Muscles were stained for 30 minutes at RT on a rotating platform using alpha-bungarotoxin (BTX) conjugated to Alexa Fluor 594 (Invitrogen) and fasciculin-2 (FCC) conjugated to Alexa fluor 488 (Invitrogen custom production, kind gift from Prof David Beeson, University of Oxford), both at 1:1,000. FDB muscles were mounted in mowiol on microscope slides and imaged using a Nikon A1R confocal system at the IMPACT Facility, University of Edinburgh. Complete, *en face* neuromuscular endplates were identified based on their BTX labelling (*70*), and BTX and FCC intensity were determined using FIJI.

## Acknowledgements

We thank Prof Norbert Polacek, University of Bern, for the careful reading of the manuscript and useful suggestions. We thank Prof David Beeson, University of Oxford, for kindly providing us with the fasciculin-II (FCC)-Alexa Fluor 488 conjugate; and the IMPACT imaging facility at the University of Edinburgh for assistance with imaging. We thank the Core Facilities Next Generation Sequencing Facility (NGS) and High Throughput Screening (HTS) at Department CIBIO University of Trento for technical support.

## Author contributions

F.L. and T.T. performed all RNA-Seq analyses. P.B. performed the SMN-specific Ribo-Seq and sub-cellular fractionation experiments. P.B. E.P. and M.C. performed all other Ribo-seq library preparation. E.G. performed all mouse tissue collection, western blotting in Figure 5 and fluorescence microscopy of NMJs. F.M. and A.I. performed the cloning of 5’UTRs for dual luciferase experiments. F.M. performed the qPCR analysis in Figure 6. M.M, M.D.S. and G.V. performed the IVTT experiments and data analysis. G.V. performed all polysomal purifications, RNA and protein extractions, western blotting and data analysis; F.L., T.T., E.G. and G.V. prepared the figures; G.V. conceived experiments and directed the research; A.Q., T.G. and G.V. obtained the funding. F.L., T.T., E.G., T.G and G.V wrote the manuscript. All authors contributed during preparation, revision and writing of the manuscript.

## SUPPLEMENTARY DATA

**Supplementary File 1**: data and mapping statistics from all RNA-Seq on brains control and at early-symptomatic SMA stage.

