## Supplementary Materials for "SMN-primed ribosomes modulate the translation of transcripts related to Spinal Muscular Atrophy"

### Supplementary Material

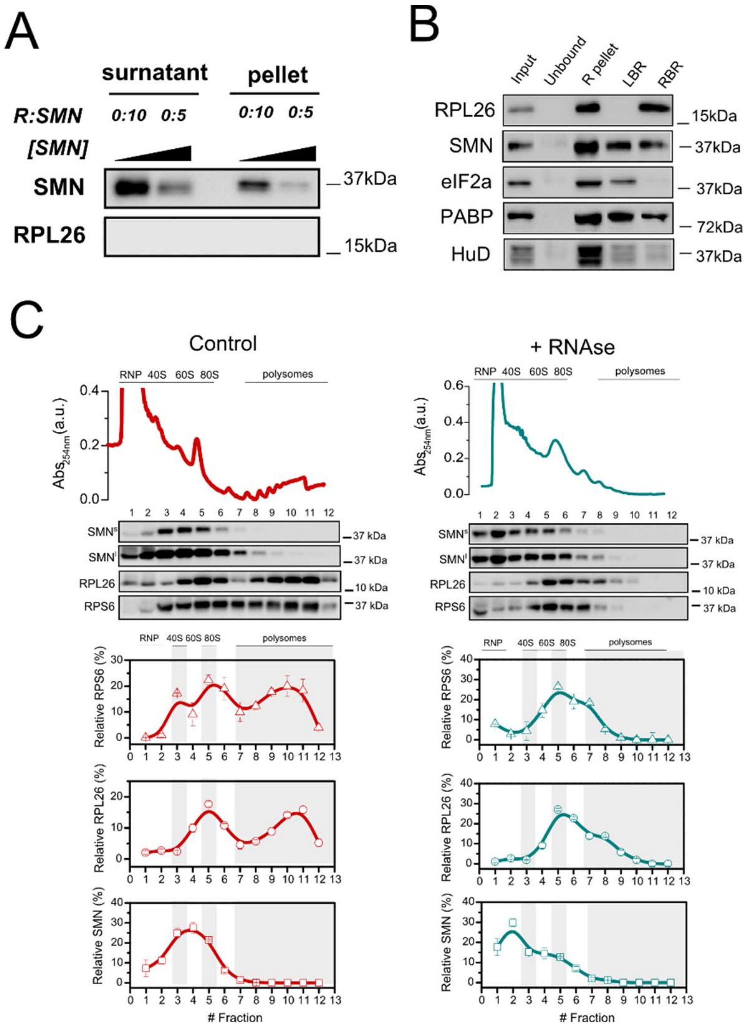


**Supplementary Figure 1**. (**A**) Determination of unspecific precipitation of recombinant SMN protein in the absence of ribosomes. (**B**) Subcellular fractionation from control P7 spinal cord and isolation of ribosomal subunits, ribosomes and polysomes (R pellet) and high salt wash analysis. Western blotting was performed on input; ribosome-free cytoplasmic components (unbound); ribosomal subunits, ribosomes and polysomes (R-pellet); loosely ribosome-bound proteins (LBR); and strongly ribosome-bound proteins (SBR). PABP and eIF2a are proteins associated to polysomes and HuD is an RNA binding protein known to be associated to polysomes through RNA interactions. The ribosomal protein L26 was used as control of ribosome sedimentation. PolyA binding protein and HuD were used as controls for RNA binding proteins interacting with polysomes though RNA dependent interactions and the elongation factor eIF2a for a loosely-associated ribosomal protein. (**C**) Polysomal profiling and co-sedimentation profiles of lysates obtained from control brain before (left panel) and after (right panel) RNAse I treatment. The ribosomal proteins RPL26 and RPS6 were used as sedimentation controls for the large and small ribosomal subunits, respectively. The sedimentation distribution of proteins along the sucrose gradient is expressed as percentage in each fraction considering as 100% the sum of all fractions. As such, only differences in the relative distribution can be observed, whilst no conclusions in absolute protein abundance differences can be drawn. The percentages shown are mean ± s.e.m. from three independent experiments.

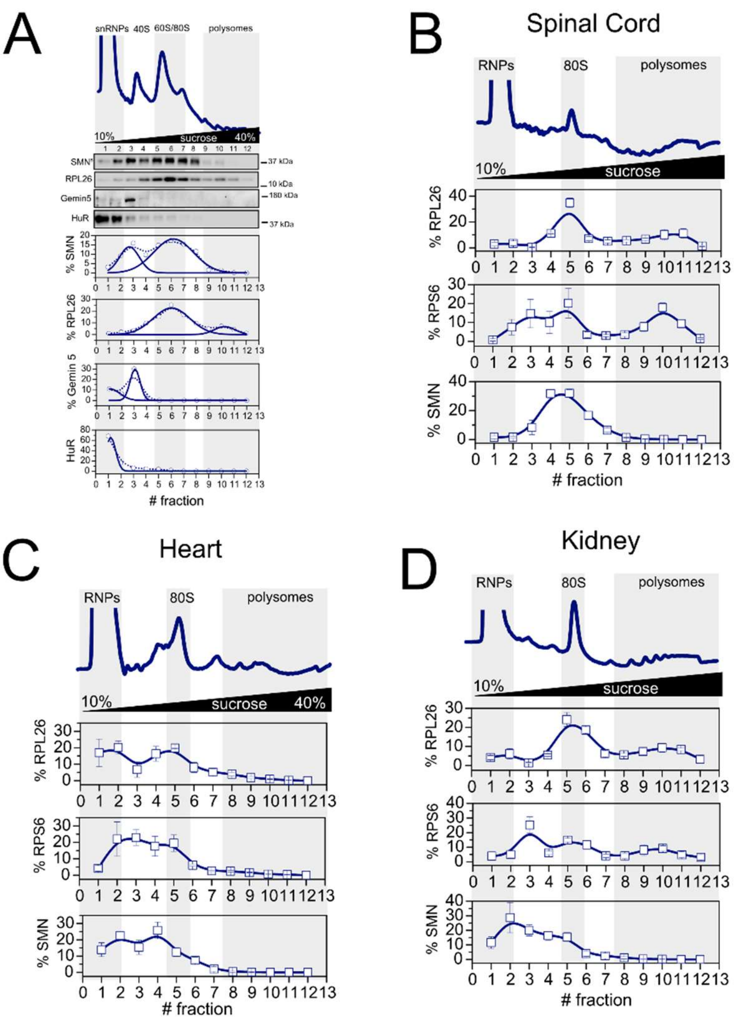


**Supplementary Figure 2**. (**A**) Co-sedimentation of SMN proteins with RiboNucleoParticles (snRNP/RNPs) and ribosomal subunits in NSC-34 lysates without cycloheximide treatment to release the ribosomal subunits. Each lane of the western blot corresponds to a fraction along the profile. The presence of two populations of SMN was established by fitting the relative co-sedimentation profile of the protein to two Gaussian curves. RPL26 was used as a marker of the 60S ribosomal subunit, Gemin 5 of the Gemin granules and HuR of RNA-granules. (**B**-**D**) Polysomal profiling and co-sedimentation profiles of lysates obtained from P3 control mouse spinal cord, heart and kidney. The ribosomal proteins RPL26 and RPS6 were used as sedimentation controls for the large and small ribosomal subunits, respectively. The sedimentation distribution of proteins along the sucrose gradient is expressed as percentage in each fraction considering as 100% the sum of all fractions. The percentage shown are mean ± sem from three independent experiments.

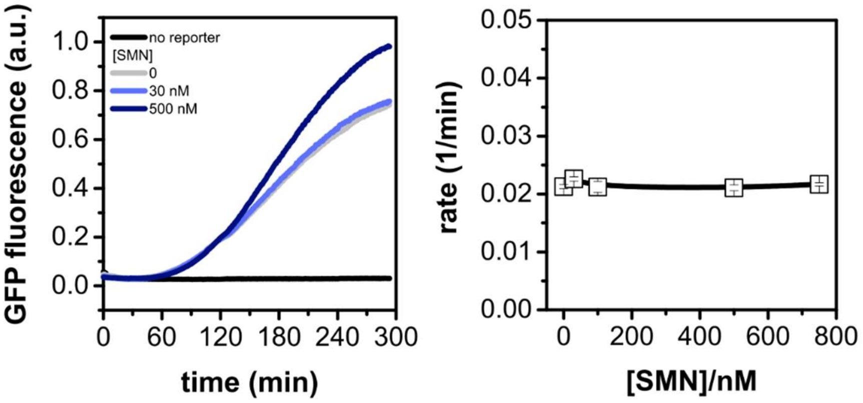


**Supplementary Figure 3**. Left panel, *in vitro* kinetic analysis of GFP translation was monitored by monitoring fluorescence increase over time after the addition of different concentrations of recombinant SMN. Right panel, comparison of the kinetic constants for different concentrations of recombinant SMN calculated from a first-order kinetics model as in Capece et al 2015 (*1*).

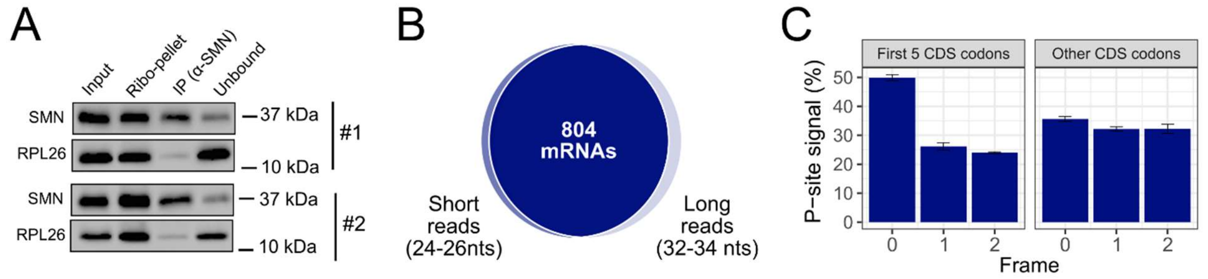


**Supplementary Figure 4**. (**A**) Two examples of immunoprecipitation of SMN from ribo-pellet in control P5 mouse brain lysates after RNAse treatment. RPL26 was used as control for immunoprecipitation of ribosomes. (**B**) Venn diagram showing the intersection of SMN-specific protein coding transcripts covered by short (24-26 nucleotides) and long (32-34 nucleotides) reads. These two populations are shared among almost all mRNAs. (**C**) Percentage of P-sites according to the three reading frames for the first five codons of the CDSs and for the remaining region of the sequences. A clear trinucleotide periodicity in the correct frame is detectable only at the beginning of the coding sequence.

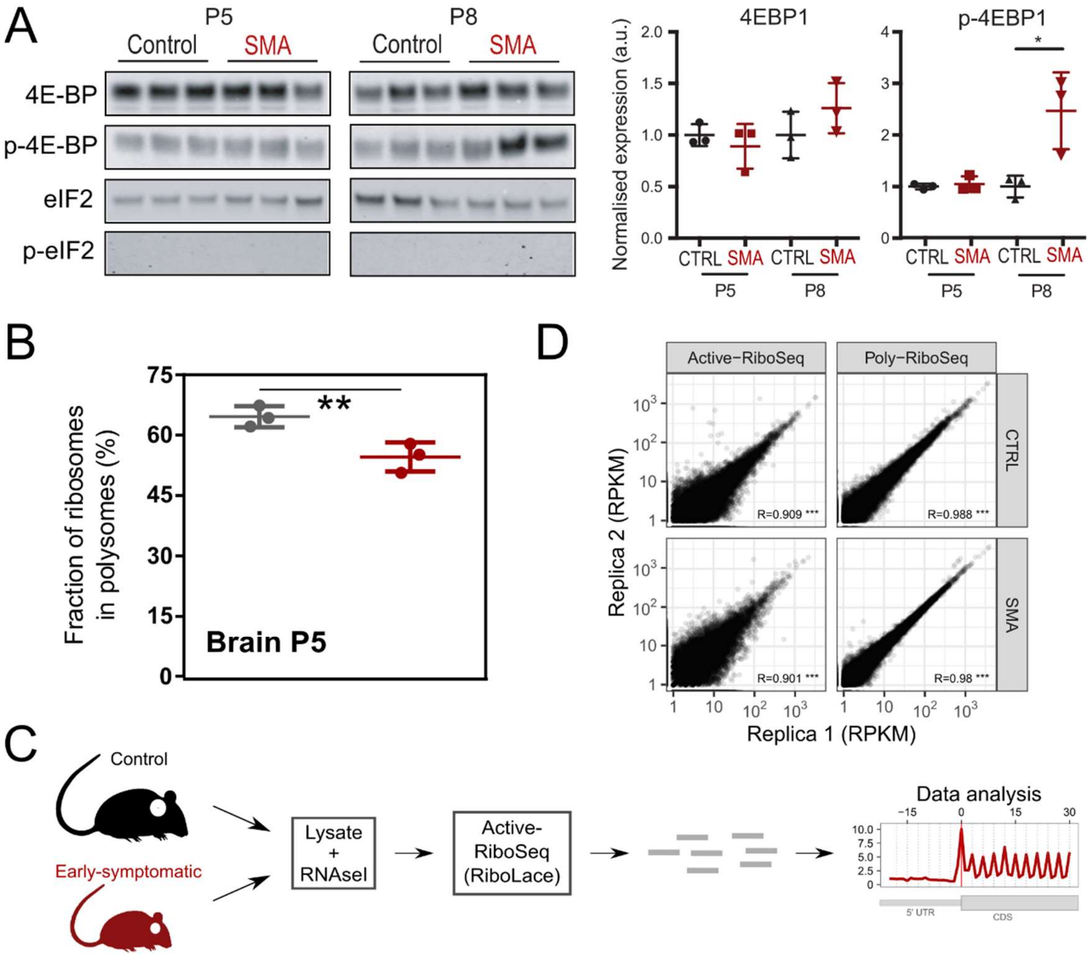


**Supplementary Figure 5**. (**A**) mTORC1 is activated only at late stages of SMA. Upper panel, total proteins from brains at early and late stages of disease (SMA) and from littermate controls (Ctrl) were probed for phosphorylation responses by immunoblotting of known mTORC1 downstream targets (4E-BP) and PERK (eIF2a). In the right panels the quantification for 4E-BP is shown (normalized to total protein stain, not shown). **p<0.01 one-way ANOVA with Tukey's multiple comparisons test, (**B**) Fraction of ribosomes in polysomes obtained from polysomal profiling of control and SMA mouse brain at early stage of disease (P5). The experiment was performed in biological triplicate from littermate mice. Significant decreases were observed by (*p < 0.05, two-tailed t-test). (**C**) Experimental design for active ribosome profiling of control and early symptomatic SMA brains. (**D**) Correlation of RPKM per protein coding transcript (49825 RNAs) between the replicas of Active-RiboSeq and Poly-RiboSeq for control and SMA samples. Correlation coefficients and statistical significance from each correlation test is shown (*** p-value < 0.001).

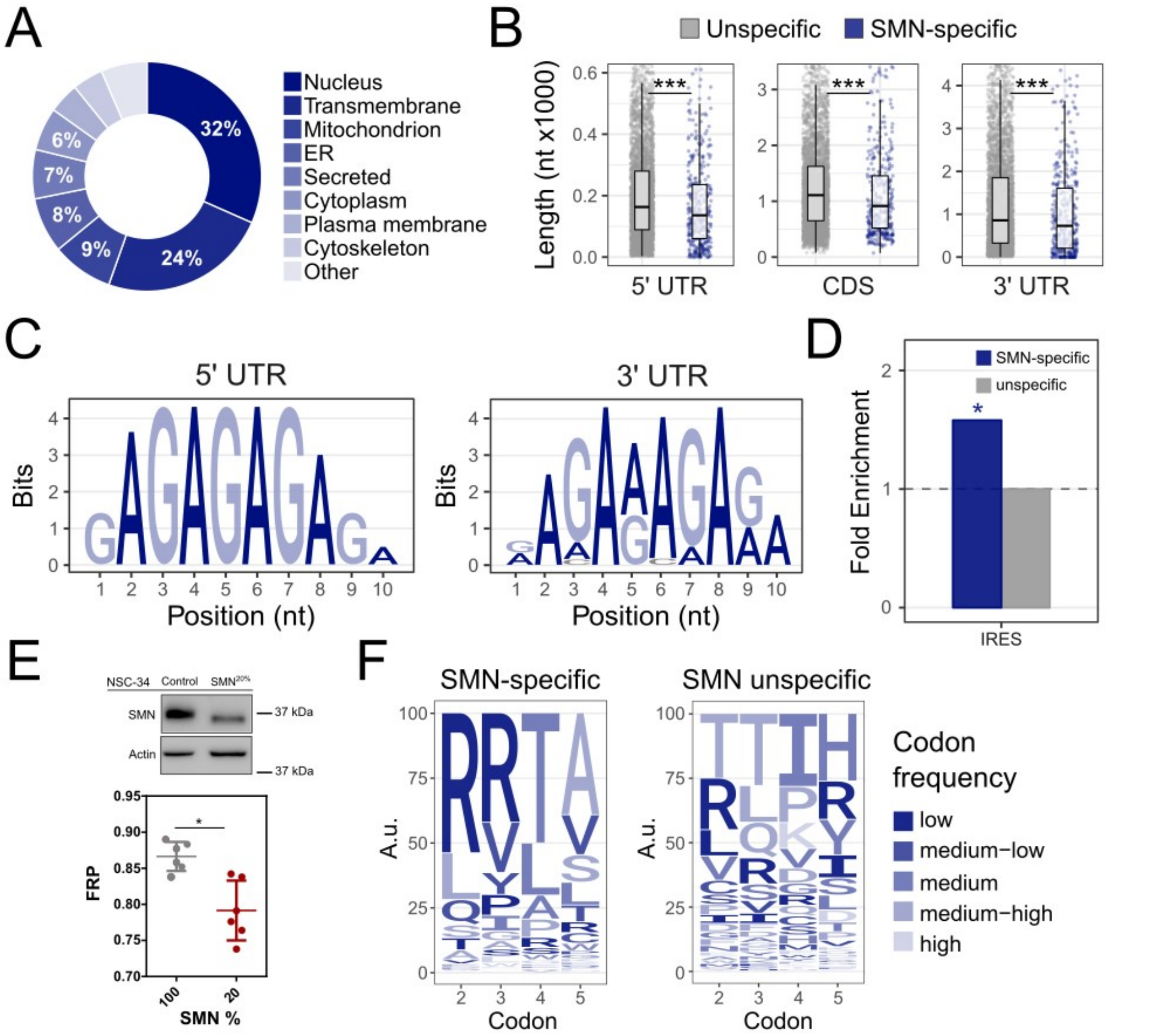


**Supplementary Figure 6.** (**A**) Identification of subcellular location of proteins synthesized by mRNAs enriched in SMN-associated ribosomes. Only manually annotated and review proteins from the UniProt database were considered. (**B**) Comparison of the lengths of coding and untranslated regions of SMN-specific transcripts, compared with unspecific transcripts selected from Poly-RiboSeq of healthy mouse brains. Statistical significance was determined with one-tailed Wilcoxon-Mann-Whitney tests. P-value: ** < 0.05, ** < 0.01, *** < 0.001. (**C**) Logo representation of top enriched motifs detected in the 5`UTR (left panel) and 3' UTR (right panel) region of SMN-specific transcripts, identified by discriminative analysis of k-mer composition. (**D**) Over-representation analysis of IRES elements among SMN-specific transcripts (in blue). IRES annotation was retrieved from Weingarten-Gabbay et al, 2016 (*2*). (**E**) SMN levels in NSC-34 cells expressing 100% SMN (control) and 20% SMN (upper panel). Comparison between the fraction of ribosomes in polysomes (FRP) in NSC-34 cells expressing 100% and 20% SMN (n = 6; t test; *p < 0.05). (**F**) Logo-like representation of the most frequent amino acids codified by SMN-specific mRNAs (left panel) and SMN unspecific RNAs (right panel) at the beginning of the coding sequence. The enrichment analysis is based on the number of occurrences of each codon, using brain-specific transcripts previously identified (see section Data Analysis of the manuscript) as background. Triplets with fold enrichments > 1 were selected and the weighted sum among synonymous codons was computed. The resulting values are displayed as percentages. Letters are colored according to the aminoacid frequency in the mouse transcriptome, divided in 5 classes (low: rare codons; high: frequent codons).

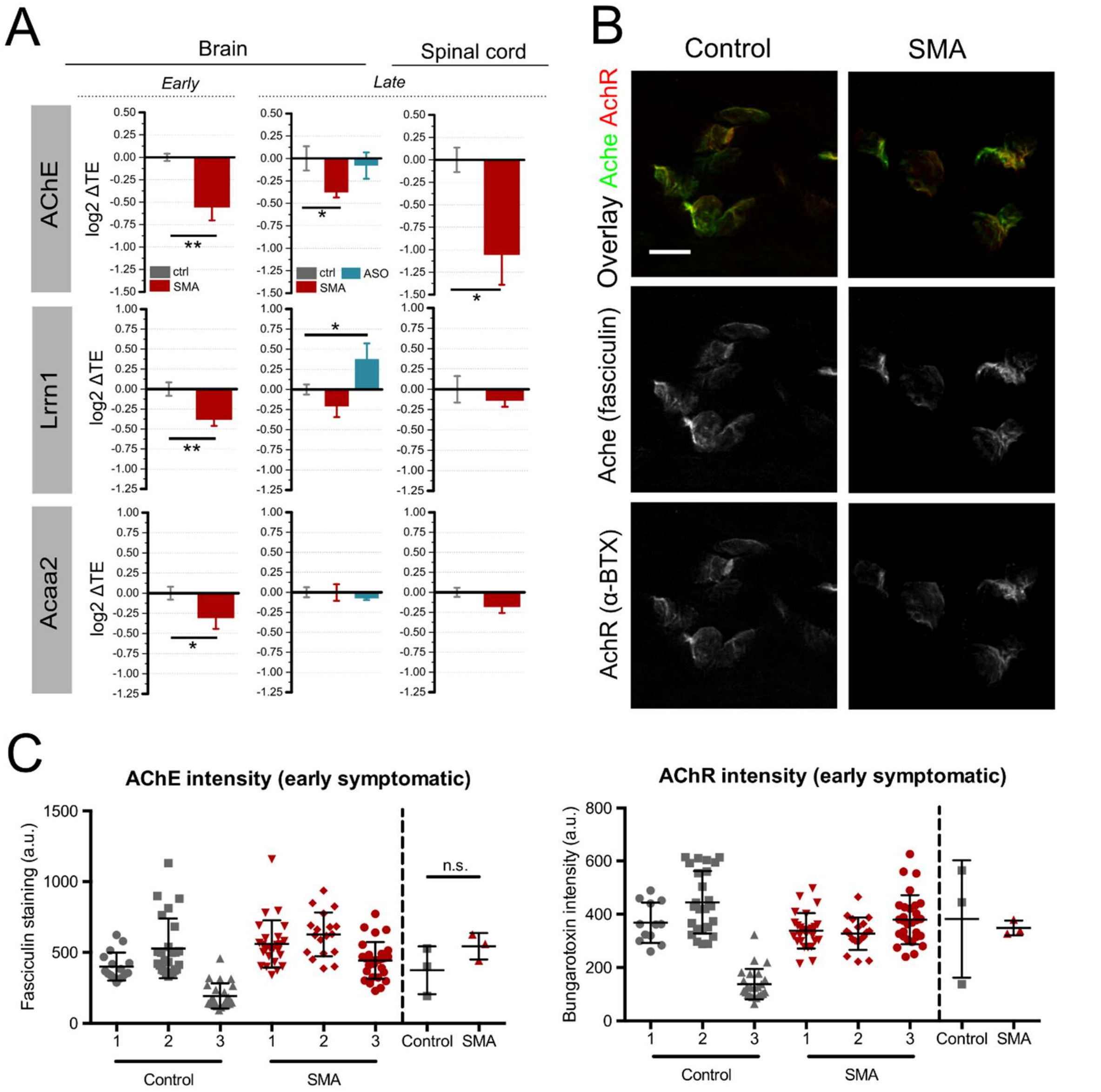


**Supplementary Figure 7**. (**A**) qPCR-derived variations of translation efficiency for three targets (Acetylcholinesterase, AChE; Lrrn1; and Acaa2) from brain at early and late symptomatic stage and in spinal cord at late stage of disease. The mean value ± SE from two to three biological replicates in 2-3 technical replicates are shown. All genes were normalized to Tuba4a (t test; *p < 0.05, **p < 0.01). (**B**) Representative images for control (left panels) and early-symptomatic SMA mouse (right panels) neuromuscular endplates. Acetylcholine receptors were labelled using alpha-bungarotoxin (BTX) conjugated to Alexa fluor 594 (red) and acetylcholinesterase was labelled using fasciculin-2 (FCC) conjugated to Alexa fluor 488 (green). Top panels show FCC / BTX overlap, middle and bottom panels show individual channels in greyscale. Scale bar: 10 μm. (**C**) FCC and BTX average intensity were determined for 2 FDB muscles in 3 control and 3 SMA mice. AChR, acetylcholine receptor; AChE, acetylcholinesterase; a.u., arbitrary units; n.s., not significant.

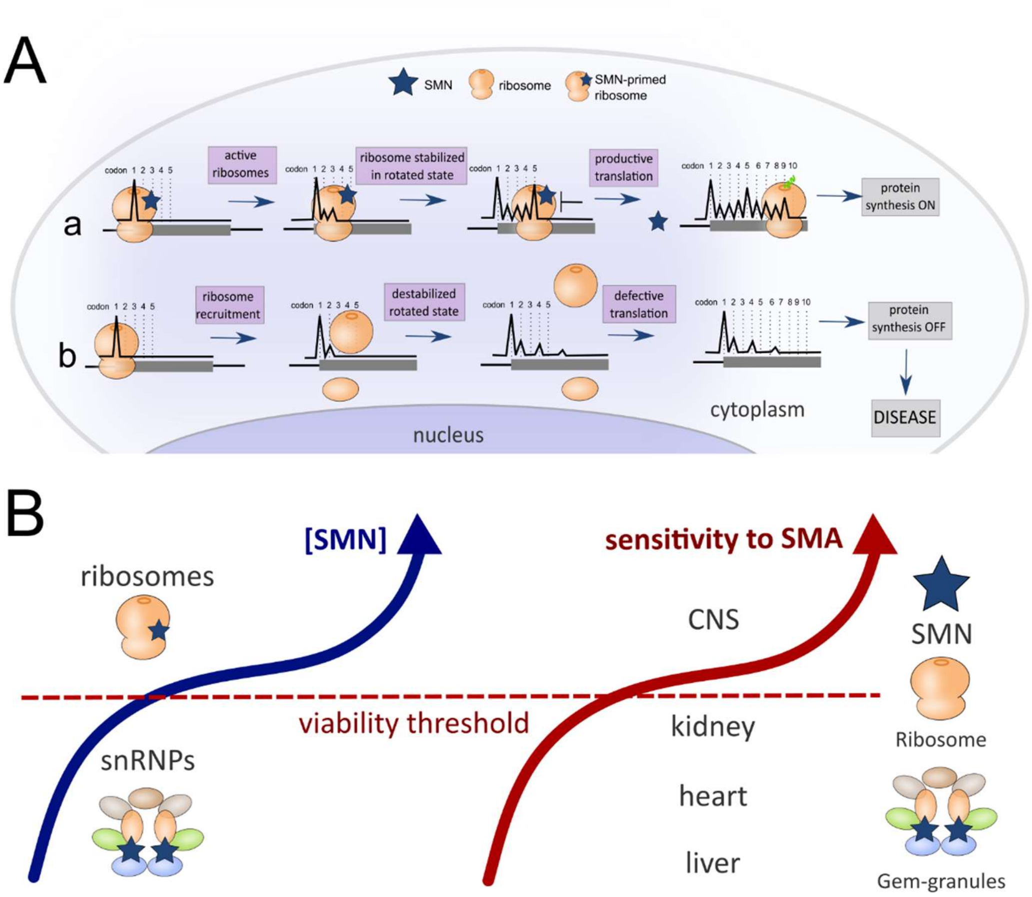


**Supplementary Figure 8**. (**A**) Schematic representation of the role of SMN-primed ribosomes in translation based on SMN-specific ribosome profiling (**Figure 4**). (a) After recruitment of ribosomes at the AUG, SMN-primed ribosomes are translating the first codons and stop at the fifth codon in a rotated state, according to the results shown in Figure 4 and to Laureau et al., 2014 (*3*). The ribosome pause at the fifth codon ensures productive translation as proposed by Han et al., 2014 (*4*) and in accordance with our results (**Figures 2** and **3**). The blue star represents SMN. (b) When SMN expression is lost in SMA, SMN-specific transcripts are bound by SMN-free ribosomes. These ribosomes cannot properly pause within the first 5 codons, causing ribosomes to drop-off (as observed by Han et al., 2014 (*4*)), leading to disruption of normal protein synthesis. (**B**) Model for tissue-specific sensitivity of tissues to decreased SMN levels in SMA and relationship to SMN-primed ribosomes. SMN binds Gemin proteins at low concentrations, while higher SMN concentrations are required for SMN to interact with ribosomes. In tissues, SMN is expressed at higher levels in the CNS (*5*) where a higher degree of SMN interaction with ribosomes/polysomes occurs. These very same tissues are also profoundly affected at the translational level by SMN loss (Bernabò et al., 2017 (*6*) and this study).

#### Supplementary Table 1. Primers for 5' UTR cloning and SMN-specific motives

| Primer | Sequence (5'- 3') |
|---|---|
| *TubA4a Fw* | AATTCTAGAATTCCGCCTCCTGCCTATAAGAGC |
| *TubA4a Rv* | AACATATGCACGTGATGAGTTTGGGCTGTACGTC |
| *AChE Fw* | AATTCTAGAATTCGGCTGTCACTGTCGGCTCAG |
| *AChE Rv* | AACATATGCACGTGGACTGCCAGGACAGGCTGGT |
| *SMN-specific Fw* | AGCTTGAATTCTATACACGTGCCACCATGCGGCGTACTCGAGC |
| *SMN-specific Rv* | CAGGGCTCGAGTACGCCGCATGGTGGCACGTGTATAGAATTCA |
| *AchE 5 aa Fw* | AGCTTGAATTCTATACACGTGCCACCATGAGGCCTCCCTGGGC |
| *AchE 5 aa Rv* | CAGGGCCCAGGGAGGCCTCATGGTGGCACGTGTATAGAATTCA |

#### Supplementary Table 2. Primers for ribosome profiling

| Primer/adaptor | Sequence (5'- 3') |
|---|---|
| *miRNA Cloning* | rAppCTGTAGGCACCATCAAT/3ddC/ |
| *RT primer* | (Phos)AGATCGGAAGAGCGTCGTGTAGGGAAAGAGTGTAGATCTCGGTGGTCGC(SpC1<br>8)-CACTCA-(SpC18)-TTCAGACGTGTGCTCTTCCGATCTATTGATGGTGCCTACAG |
| Forward library | AATGATACGGCGACCACCGAGATCTACAC |
| *Index 1* | CAAGCAGAAGACGGCATACGAGATCGTGATGTGACTGGAGTTCAGACGTGTGCTCTTCCG |
| *Index 2* | CAAGCAGAAGACGGCATACGAGATACATCGGTGACTGGAGTTCAGACGTGTGCTCTTCCG |
| *Index 3* | CAAGCAGAAGACGGCATACGAGATGCCTAAGTGACTGGAGTTCAGACGTGTGCTCTTCCG |
| *Index 4* | CAAGCAGAAGACGGCATACGAGATTGGTCAGTGACTGGAGTTCAGACGTGTGCTCTTCCG |
| *Index 5* | CAAGCAGAAGACGGCATACGAGATCACTGTGTGACTGGAGTTCAGACGTGTGCTCTTCCG |
| *Index 6* | CAAGCAGAAGACGGCATACGAGATATTGGCGTGACTGGAGTTCAGACGTGTGCTCTTCCG |
| *Index 7* | CAAGCAGAAGACGGCATACGAGATGATCTGGTGACTGGAGTTCAGACGTGTGCTCTTCCG |
| *Index 8* | CAAGCAGAAGACGGCATACGAGATTCAAGTGTGACTGGAGTTCAGACGTGTGCTCTTCCG |
| *Index 9* | CAAGCAGAAGACGGCATACGAGATCTGATCGTGACTGGAGTTCAGACGTGTGCTCTTCCG |
| *Index 10* | CAAGCAGAAGACGGCATACGAGATAAGCTAGTGACTGGAGTTCAGACGTGTGCTCTTCCG |
| *Index 11* | CAAGCAGAAGACGGCATACGAGATGTAGCCGTGACTGGAGTTCAGACGTGTGCTCTTCCG |
| *Index 12* | CAAGCAGAAGACGGCATACGAGATTACAAGGTGACTGGAGTTCAGACGTGTGCTCTTCCG |

#### Supplementary Table 3. Primers for ddPCR and RT-qPCR

| Gene | Primer | Sequence (5'- 3') |
|---|---|---|
| *AChE* | Forward | CGCTTTCTCCCCAAATTGCT |
| | Reverse | GTTCTTCCAGTGCACCATGT |
| *TubA4a* | Forward | AGGAAGTAGGCATCGACTCC |
| | Reverse | AACACAGTGAACAGGCTCCA |
| *Acaa2* | Forward | CAGGAGACCTTCGAGTCAGC |
| | Reverse | CTCCGATGGTTGGTGTTTTT |
| *Lrrn1* | Forward | TGTGCTTGACCCCTTTCTTC |
| | Reverse | AATGCTGCTTCCAAACCCTA |
